# Bayesian evidence for the neural dissociation between finger and hand imitation skills

**DOI:** 10.1101/2024.01.23.576674

**Authors:** Hannah Rosenzopf, Lisa Röhrig, Georg Goldenberg, Hans-Otto Karnath

## Abstract

**Introduction:** For limb apraxia ‒ a heterogeneous disorder of higher motor cognition following stroke ‒ an enduring debate has arisen regarding the existence of dissociating neural correlates for finger and hand gestures in the left hemisphere. We re-assessed this question asking whether previous attempts analysing pooled samples of patients with deficits in only one and patients with deficits in both imitation types might have led to systematically biased results.

**Methods:** We conducted frequentist and Bayesian, voxelwise and regionwise lesion symptom mappings on a pooled sample (N=96) and subsamples containing only shared and only isolated hand and finger imitation deficits and respective controls.

**Results:** Anatomical analyses on the isolated sample reinforced a cortical dissociation of finger deficits (located more anteriorly) and hand deficits (located more posteriorly). The presence of patients with shared deficits did indeed dilute associations that appeared stronger in the respective isolated samples. Also, brain regions truly associated with hand imitation deficits showed a positive bias for finger imitation deficits, when the sample contained patients with shared deficits. In addition, our frequentist parameters uncovered that some of our Bayesian evidence supported reverse associations (damage protecting from rather than increasing the deficit).

**Discussion:** Joint analyses of patients with shared and isolated imitation deficits do indeed lead to biases, which may explain why some previous studies have failed to detect the actual neural dissociation between hand and finger imitation deficits.

## Introduction

Limb apraxia is a heterogeneous disorder of higher motor cognition, impairing the execution of different types of actions (Randerath, 2023). The earliest research distinguished between ideational, ideomotor, and limb kinetic apraxia (Liepman, 1920). In an attempt to create a more symptom-oriented classification, Goldenberg (2011), introduced a subdivision into the apraxias of (1) tool use, (2) gesture production, and (3) imitation. The apraxia of imitation is further subdivided based on the affected body part, i.e. the hand (De Renzi et al., 1980; Goldenberg, 1996; Goldenberg & Karnath, 2006; Kimura & Archibald, 1974; Lehmkuhl et al., 1982), fingers (Goldenberg, 1996; Goldenberg & Karnath, 2006; Goldenberg & Strauss, 2002), legs (Lehmkuhl et al., 1982), feet (Goldenberg & Strauss, 2002), as well as different facial features (Bizzozero et al., 2000; Lehmkuhl et al., 1982).

While hemispheric differences based on the impaired body part support behavioral dissociations (Bizzozero et al., 2000; Goldenberg, 1996; Goldenberg & Strauss, 2002; Lausberg & Cruz, 2004), a lasting controversy about whether or not finger and hand gestures have dissociating neural correlates in the left hemisphere has emerged. Reports of dissociating neural correlates for finger and hand imitation deficits in the left hemisphere (Goldenberg, 1996; Goldenberg & Karnath, 2006; Haaland et al., 2000) seemingly contradict more recent accounts, which failed to uncover such dissociation (Achilles et al., 2017; Hoeren et al., 2014). In particular, Achilles and colleagues (2017) aimed at eliminating this ambiguity once and for all by applying Bayesian statistics, which they judge as particular beneficial in this context, due to its ability to uncover evidence for both alternative and null hypothesis. Their results, derived from a large sample of 257 left hemisphere stroke patients, discount previous evidence for the dissociation, arguing that it had been reported only by small samples that had not performed any statistical evaluations. However, another crucial difference between the latter studies demonstrating dissociating neural substrates and those that did not, including the one by Achilles and colleagues (2017), is a fundamentally different sampling approach. While the earlier descriptive studies (that found a dissociation) exclusively compared patients with isolated hand and finger imitation deficits to controls without the respective deficit, the more recent statistical analyses (that did not find a dissociation) assigned patients with both shared and isolated deficits to the same group.

Grouping patients with shared and isolated imitation deficits together could potentially be problematic, in particular when investigating the existence of a task dissociation. We know that the lesions of patients with an isolated deficit can in theory only be associated with that deficit but not the other (which in itself should already be considered as support for a dissociation). Lesions of patients with a shared deficit of hand and finger imitation skills, could contain what henceforth will be considered *shared areas* (i.e. areas whose damage leads to a simultaneous deficit in hand and finger imitation skills), or will at least contain voxels of both kinds of *isolated areas* (i.e. areas leading to only hand or only finger imitation deficits). In scenario one, a voxel truly causing an isolated deficit will reduce the power of voxels truly associated with a shared deficit and vice-versa. In scenario two, patients with a shared deficit will have simultaneous damage in *isolated hand* - and *isolated finger areas*. That is, voxels leading to the respective other deficit will unavoidably be represented in patients’ lesions, therefore potentially introducing a systematic bias. Both scenarios could lead to the wrongful rejection of a truly existing dissociation, the former by uncovering only *shared areas* but not the underpowered isolated ones, the latter by erroneously associating *isolated areas* for one deficit also with the respective other, due to their frequent co-occurrence in patients with a shared deficit.

Achilles and colleagues (2017) assigned damage to brain regions associated with imitation skills a higher susceptibility to hand gesture deficits and argued that finger deficits might accompany particularly pronounced problems with hand imitation. At the same time, the authors reported ten patients with the opposite dissociation (i.e. patients with an isolated finger imitation deficit). They argued that eight of those cases, who had a very mild isolated finger imitation deficit might be considered noise by the model but admit that their rationale does not sufficiently explain the remaining two cases with a severe isolated finger imitation deficit. In light of the methodological dependencies described above, a potential alternative explanation could be that the results obtained by Achilles and colleagues (2017) reflect the larger subgroups with shared deficits, which might indeed have a higher susceptibility for hand than for finger gestures. The higher frequency of *shared areas* could have outpowered *isolated areas.* Alternatively, isolated hand areas could have been wrongly associated to finger imitation deficits.

In the present study, we investigated whether appropriate subsampling can indeed uncover dissociating neural correlates of hand and finger imitation deficits that are masked up in a joint analysis of patients with dissociating and shared imitation deficits. We recreated the analyses on hand and finger imitation scores reported by Achilles and colleagues (2017) with some adaptations. First, in addition to analyses on the full sample we also created sub-samples. In one sub-sample, patients with the respective isolated deficit were excluded. This sample therefore represented only patients with the deficit in question, if it co-occurred with the respective other imitation deficit (reduced shared sample). In the other sub-sample, all patients with a shared deficit were excluded; it thus included only those patients in whom the deficit of interest occurred in isolation from the respective other imitation deficit (reduced isolated sample). Second, we applied both frequentist and Bayesian approaches on both the voxel and the region level. First, we hypothesised that the reduced isolated sample would uncover dissociating voxels and brain regions with higher statistical parameters than in the (negatively biased) full samples. Second, we, conversely, expect brain areas in the full sample to show a positive bias for those brain regions truly associated with the respective other imitation deficit due to the presence of those lesion sites in patients with a shared deficit. The second hypothesis is limited to regionwise analyses since the reduced dimensionality of the regionwise approach offers the possibility – in contrast to the voxelwise approach – to demonstrate the effects of the different sampling approaches on specific data points throughout the different sampling conditions. Thirdly, shared areas (if they exist) should have the best chance of being uncovered in the reduced shared sample in both analyses on hand and finger imitations scores but should not be found in analyses on the isolated samples.

## Methods

### Participants

Data of 96 brain-damaged patients investigated at the Neuropsychological Department of the Bogenhausen Hospital in Munich, Germany, were analysed retrospectively. The identical sample has previously been used to investigate associations between apraxia and aphasia (Goldenberg & Randerath, 2015). It includes 23 female and 73 male patients who had suffered a first-time left hemispheric stroke on average 13.3 weeks (range 3 - 58) before the behavioural examination. 73 patients had suffered an ischemic stroke, 23 a haemorrhagic infarct. Additional demographic and clinical data, divided by patient sub-groups (see below), can be derived from table 1. Consent for the scientific reuse of their data was provided by the patients; We followed ethical standards as predefined by the revised Declaration of Helsinki.

**Table 1:**
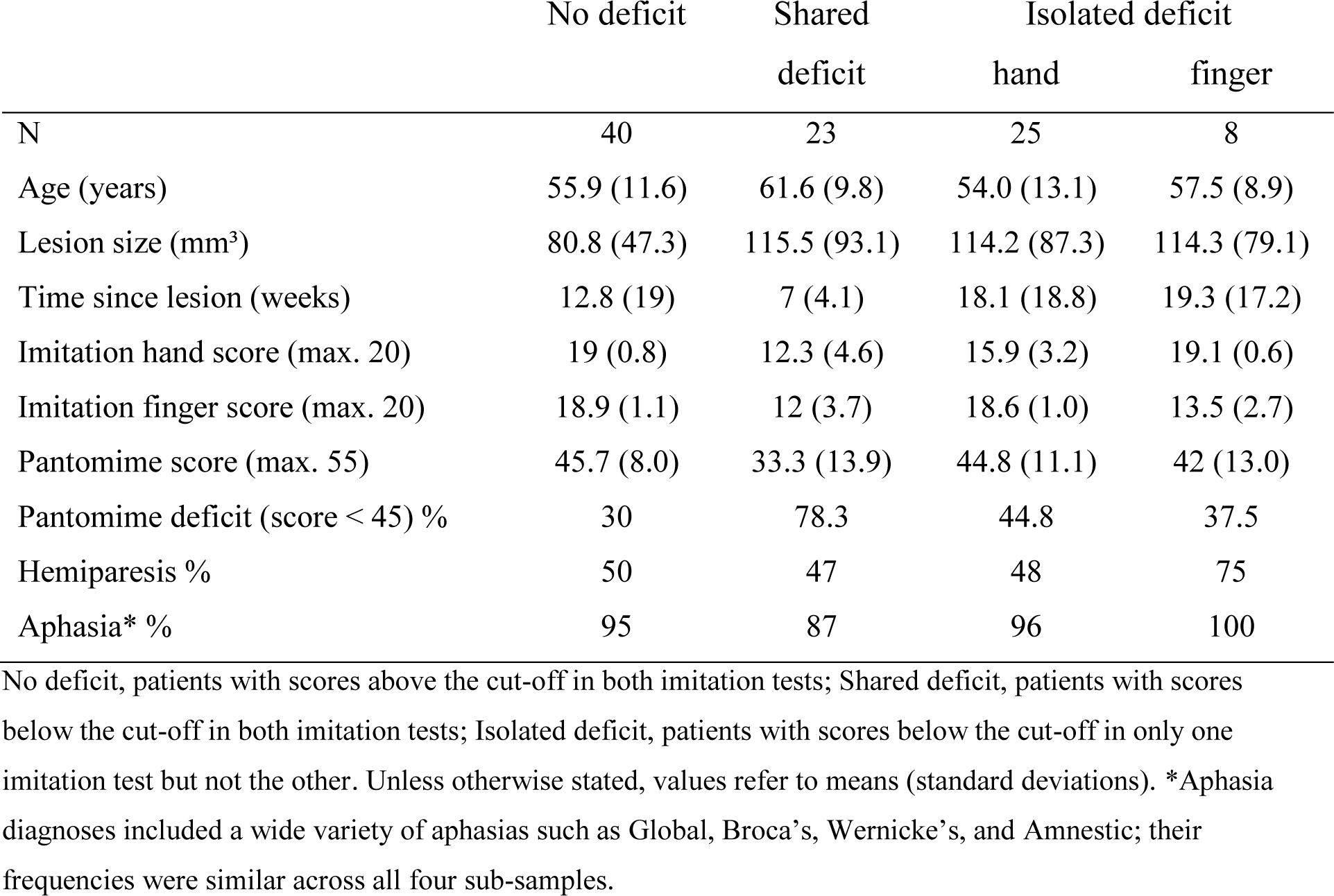
Descriptive statistics concerning relevant demographical and clinical data divided by patient sub-group.

|  | No deficit | Shared deficit | Isolated deficit |  |
| --- | --- | --- | --- | --- |
|  |  |  | hand | finger |
| N | 40 | 23 | 25 | 8 |
| Age (years) | 55.9 (11.6) | 61.6 (9.8) | 54.0 (13.1) | 57.5 (8.9) |
| Lesion size (mm<sup>3</sup>) | 80.8 (47.3) | 115.5 (93.1) | 114.2 (87.3) | 114.3 (79.1) |
| Time since lesion (weeks) | 12.8 (19) | 7 (4.1) | 18.1 (18.8) | 19.3 (17.2) |
| Imitation hand score (max. 20) | 19 (0.8) | 12.3 (4.6) | 15.9 (3.2) | 19.1 (0.6) |
| Imitation finger score (max. 20) | 18.9 (1.1) | 12 (3.7) | 18.6 (1.0) | 13.5 (2.7) |
| Pantomime score (max. 55) | 45.7 (8.0) | 33.3 (13.9) | 44.8 (11.1) | 42 (13.0) |
| Pantomime deficit (score < 45) % | 30 | 78.3 | 44.8 | 37.5 |
| Hemiparesis % | 50 | 47 | 48 | 75 |
| Aphasia* % | 95 | 87 | 96 | 100 |

No deficit, patients with scores above the cut-off in both imitation tests; Shared deficit, patients with scores below the cut-off in both imitation tests; Isolated deficit, patients with scores below the cut-off in only one imitation test but not the other. Unless otherwise stated, values refer to means (standard deviations). \*Aphasia diagnoses included a wide variety of aphasias such as Global, Broca's, Wernicke's, and Amnesic; their frequencies were similar across all four sub-samples.

### Behavioural testing and sub sampling

Diagnostic scores concerning the apraxia of the imitation of hand and finger gestures (Goldenberg, 1996) served as the two dependent variables in our analyses. For both tests, ten hand or finger postures, are modelled by the investigator. After having demonstrated a gesture, the investigator returns to a neutral posture before asking the patient to imitate the previously seen gesture. Two points are scored for a correct imitation at the first attempt. If the patient successfully imitated the gesture at the second attempt after seeing the gesture a second time, they received one point. This results in a maximum of 20 points for each test. The cut-off for pathological performance in the hand imitation test is <18, the cut-off for pathological performance for the finger imitation test is <17 (Goldenberg, 1996). Other noteworthy assessments included the Aachen Aphasia Test (ATT) and a 20-item test assessing the apraxia of pantomime (Goldenberg, 2003; Goldenberg et al., 2007). In addition to the full sample (N=96), which contained all patients, different sub-samples representing different imitation deficit patterns were created. The reduced shared hand imitation sample (N=71) did not contain the patient group with an isolated hand imitation deficit and the reduced shared finger imitation sample (N=88) did not contain patients with an isolated finger imitation deficit. Patients with an isolated deficit of the respective other type, here served as part of the respective control groups. The former two groups represented patients with shared hand- and finger imitation deficits, respectively. Patients with a shared imitation deficit were excluded from the reduced isolated sample (N=72). The reduced isolated sample (like the full sample) was therefore the same for hand and finger imitation.

### Imaging and Image processing

CT (N=12) and MRI (N=84) imaging obtained within three weeks from the behavioural testing was used to manually map patients’ lesions on axial slices of a T1-weighted template in MNI space, which is accessible via MRIcro software (Rorden & Brett, 2000; http://people.cas.sc.edu/rorden/mricro/index.html). Eleven slices incrementing in steps of eight millimeter from z-coordinate −40 to 40 plus a twelfth one representing z-coordinate 50 were used to map patients’ lesions to the respective (or the closest) available axial slice. These slices were 3-dimensionally reconstructed using a MATLAB script previously used for this purpose (Klingbeil et al., 2020). This script expands the existing slices along the z-axis and smooths the resulting images with a Gaussian kernel. The lesions were then (re)binarized at a threshold of 0.45 (for details see Klingbeil et al., 2020). Lesion overlays divided by the four patient sub-groups (see above) are illustrated in figure 1.

**Figure 1:**
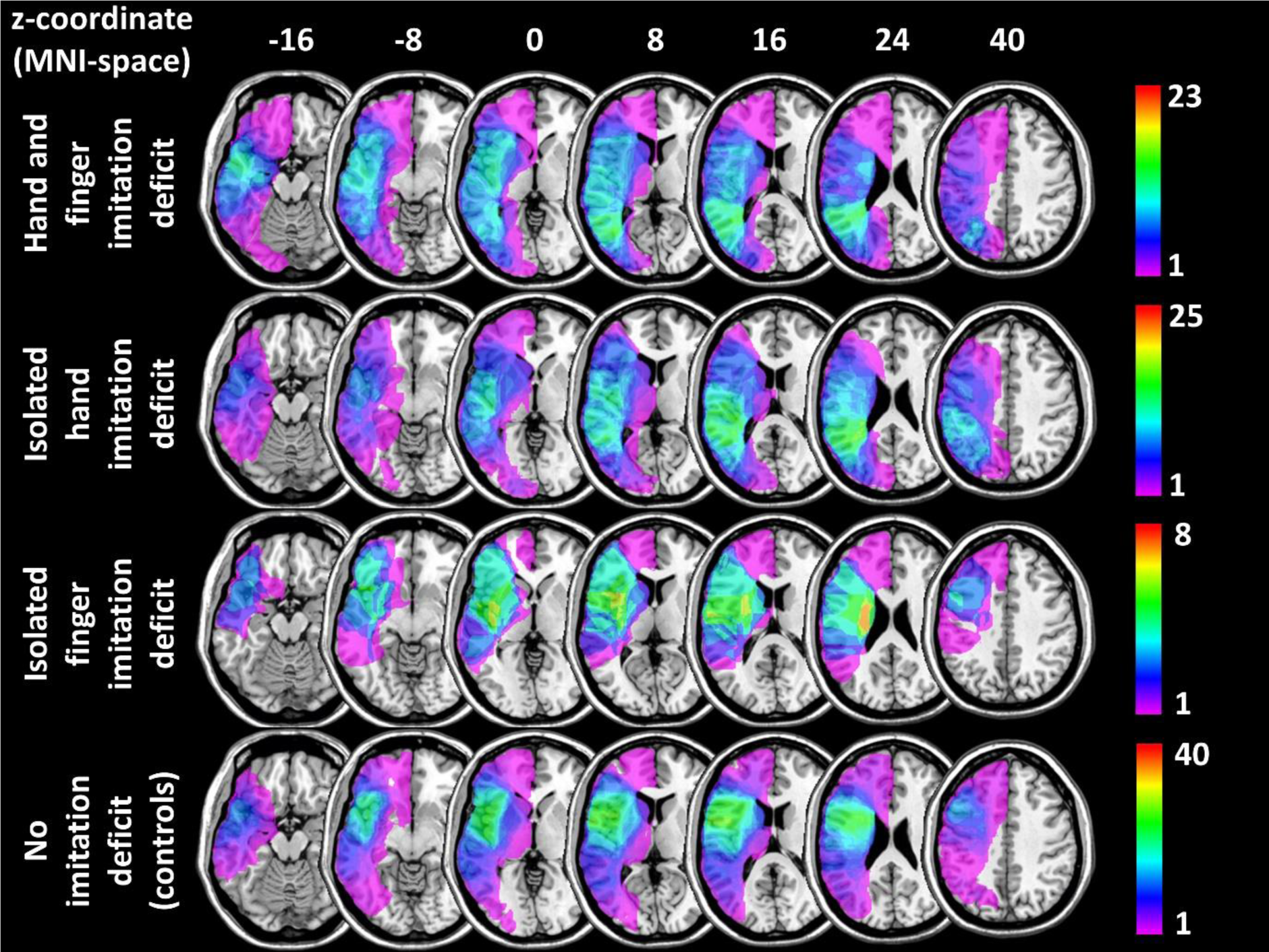
Lesion overlays for different patient groups.

### Data analysis

Depending on the imitation type investigated, voxelwise as well as regionwise brain-behaviour associations were modelled using either the hand or the finger imitation score as the dependent variable. Voxelwise lesion-symptom mapping (VLSM) was applied based on frequentist mass-univariate General Linear Models (GLMs) using NiiStat (https://github.com/neurolabusc/NiiStat) and Bayesian GLMs using the Bayesian lesion-deficit inference (BLDI) software (Sperber et al., 2023). Both approaches were set to include voxels damaged in at least 5 patients. Results of the frequentist approach were corrected for multiple comparisons using False Discovery Rate at a corrected p < .01 (FDR; Benjamini & Hochberg, 1995). The statistical parameters returned by NiiStat are z-scores. To keep results comparable to those in the study by Achilles and colleagues (2017), we adopted their definition of strong evidence as a Bayes Factor (BF) > 20. Unlike the BLDI, which uses the 10^th^ logarithm, Achilles and colleagues (2017) used the natural logarithm, so we adapted the respective line in the BLDI script. In the applied version of the logBF, zero marks the border between evidence for H1 (represented by positive numbers) and evidence for H0 (represented by negative numbers), the equivalent to our cut-off (BF>20) is a logBF>2.99. VLSMs were conducted both with and without controlling for lesion size. The effects in Bayesian frameworks are reported in the main text, detailed results from the frequentist approaches can be found in supplementary figures 1-4. Patient’s regionwise lesion load was assessed using custom scripts created using MATLAB 2019b, which overlapped each patient’s lesion with all left hemispheric parcels derived from the Desikan-Killiany atlas (Desikan et al., 2006) provided by the IIT Human Brain Atlas (v.5.0) (https://www.nitrc.org/projects/iit/). Anatomical terms throughout the manuscript follow the atlas and its labels. For the frequentist GLMs, we adapted customized MATLAB scripts, previously used for similar purposes (Röhrig et al., 2023). Correction for multiple comparisons was based on maximum-statistics permutation. 5000 pseudo-random permutations were performed, results are again reported for a corrected p < .01. In contrast to NiiStat, our MATLAB scripts return t-scores instead of z-scores. While the directionality here is the opposite (i.e. positive t-scores indicate the expected association between damage and worse performance) both are frequentist estimates of the associations and their directions. Since no script for a regionwise Bayesian GLM existed within the BLDI software, we customized the BLDI disconnection script to serve this purpose. Note that results, in the main text are reported in BFs to avoid the underrepresentation of higher evidence, while logBFs were used for figures depicting VLSM results, to enhance the differentiability of the colour gradients.

## Results

### Voxelwise lesion-symptom mapping

#### Imitation of hand gestures

Applying the Bayesian GLM on the full sample to uncover in the neural correlates of hand imitation skills led to 23842 voxels (BF_max_=513,733.53). The same analysis on the reduced shared deficit sample led to 5639 voxels (BF_max_= 24,631.15). In the reduced isolated deficit sample 39442 voxels (BF_max_= 5,928,694.17) were observed. See figure 2 for voxels with sufficient evidence for an association between voxel and hand imitation deficit. All analyses uncovered a larger cluster around the borders of the occipital parietal and temporal lobe. Beyond, analyses on the full and the reduced shared hand sample showed a smaller frontal cluster. Figure 3 illustrates discrepancies between analyses with and without control for lesion size. We observed that the application of lesion size control overall increased the number of both anterior and posterior voxels. Interestingly, the anterior cluster did not only resurface here, it even formed the largest anterior cluster among the three sub-samples.

**Figure 2:**
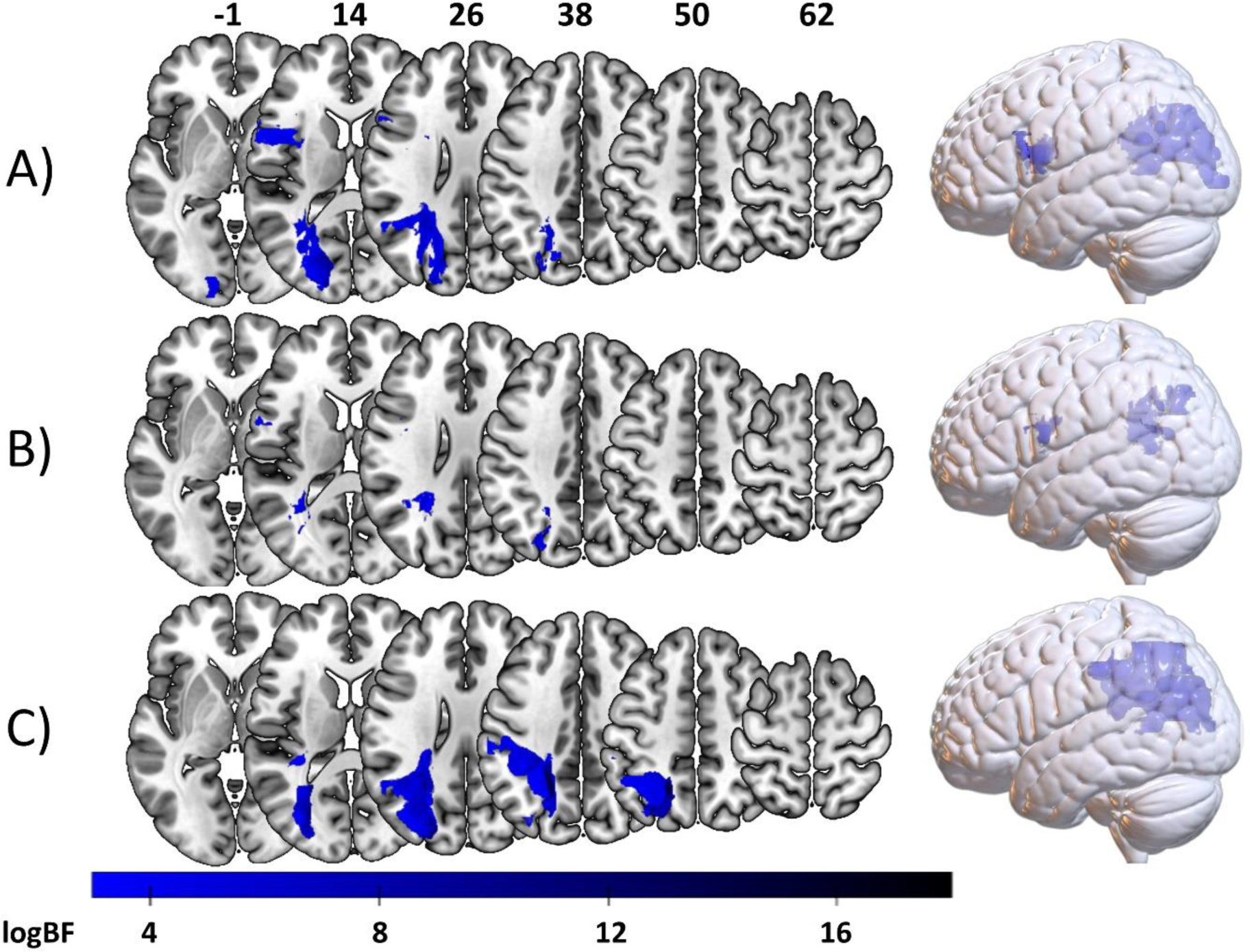
Bayesian associations between voxelwise brain damage and hand imitation deficits. Depicted are all voxels with a logBF > 2.99. Voxels associated with hand imitation deficits based on the VLSM on A) the full sample, B) the shared deficit sample, and C) the isolated sample.

**Figure 3:**
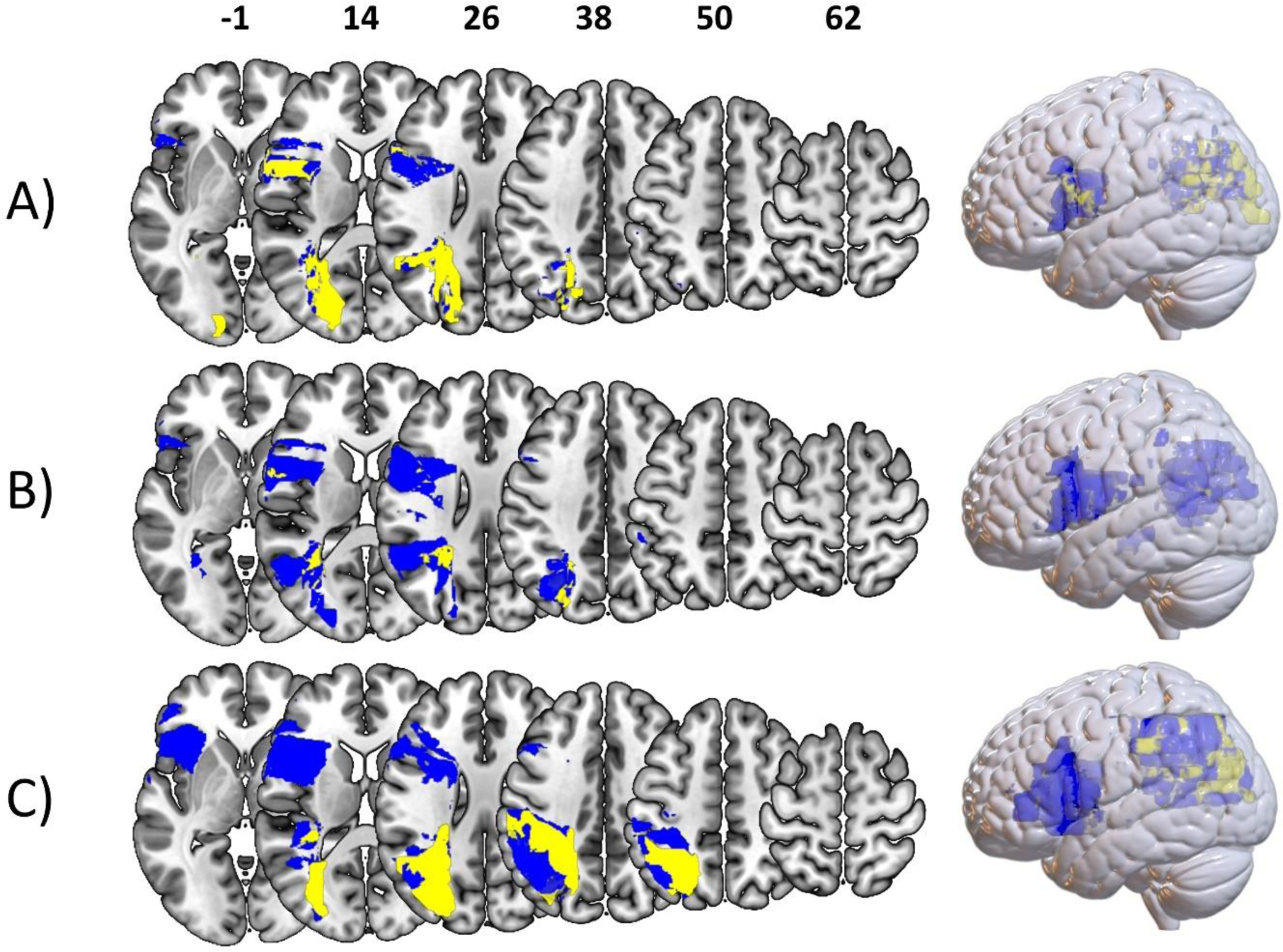
Comparison of results derived from the same analyses with vs. without lesion size control on hand imitation scores. Blue represents voxels with sufficient evidence for the H1 with lesion size control only, there were no voxels with sufficient evidence for the H1 without lesion size control only. Yellow voxels mark voxels uncovered by both analyses. A) Displays results from analyses on the full sample. B) Showcases voxels derived from analyses on the reduced shared sample. C) Represents voxels uncovered in the reduced isolated sample.

#### Imitation of finger gestures

In the full sample, the BLDI uncovered small anterior voxel clusters associated with the finger imitation score, containing 321 voxels in total (BF_max_= 3127.50). For the reduced shared deficit sample, 299 voxels in total (BF_max_= 2544.70) were found. They formed minimal clusters both anteriorly and more posterior. In the reduced isolated deficit sample, the Bayesian approach led to considerably larger voxel clusters containing 11373 voxels in total (BF_max_=8,780,833.98). with a strong anterior focus in the pars orbitalis and the rostral divison of the middle frontal gyrus (rMFG). Voxel clusters are visualized in figure 4. Figure 5 offers an overview of differences between the results reported here and those including lesion size control. We observed only minimal voxel clusters in the full and reduced shared sample, regardless of lesion size control. In the reduced isolated sample, the analysis without lesion size control yielded overall more significant voxels. A large part of the anterior cluster, however, was confirmed by both analyses.

**Figure 4:**
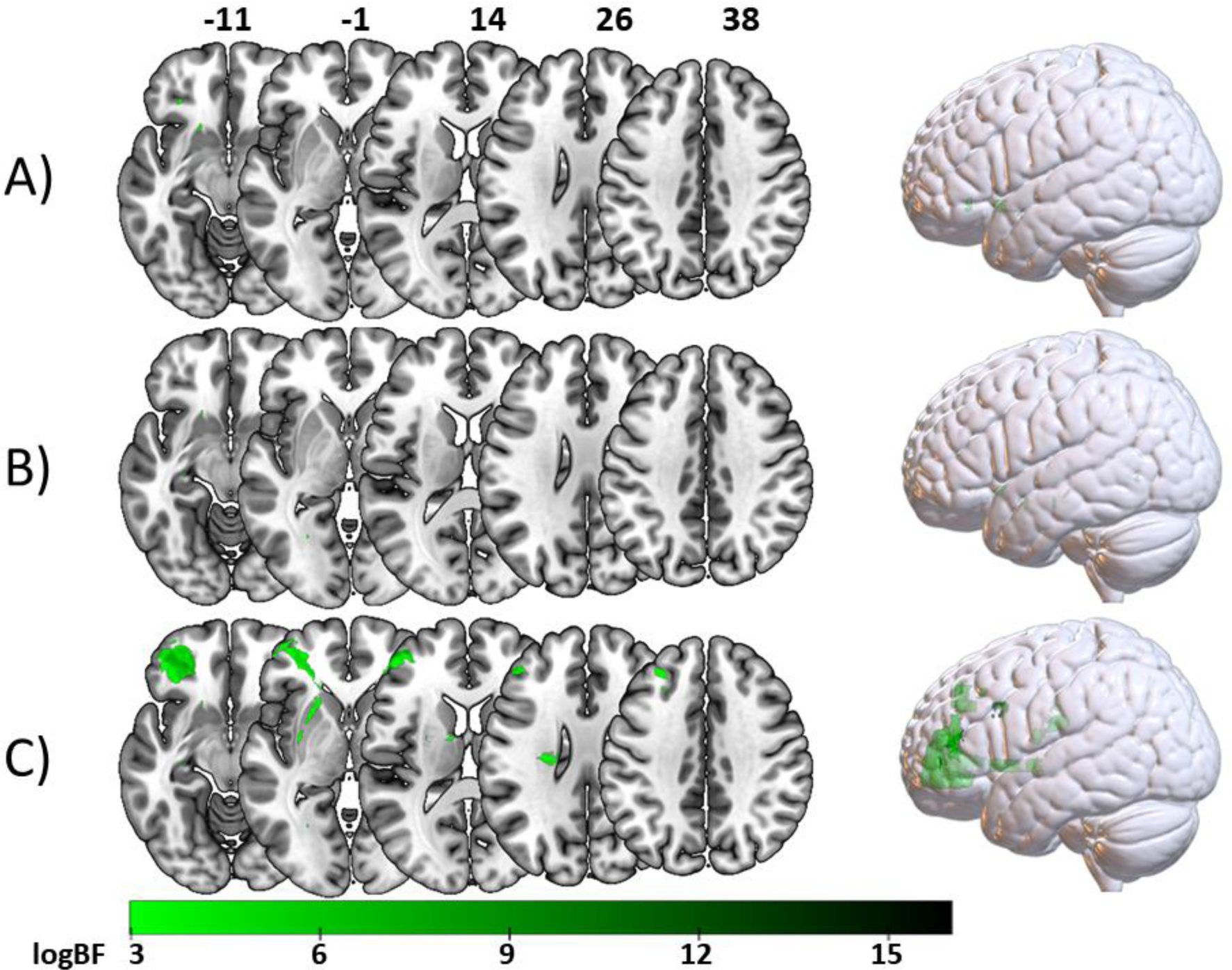
Bayesian associations between voxelwise brain damage and finger imitation deficits. Depicted are all voxels with a logBF > 2.99. Voxels resulting from the VLSM on A) the full sample, B) the reduced shared finger deficit sample, and C) the reduced isolated finger sample.

**Figure 5:**
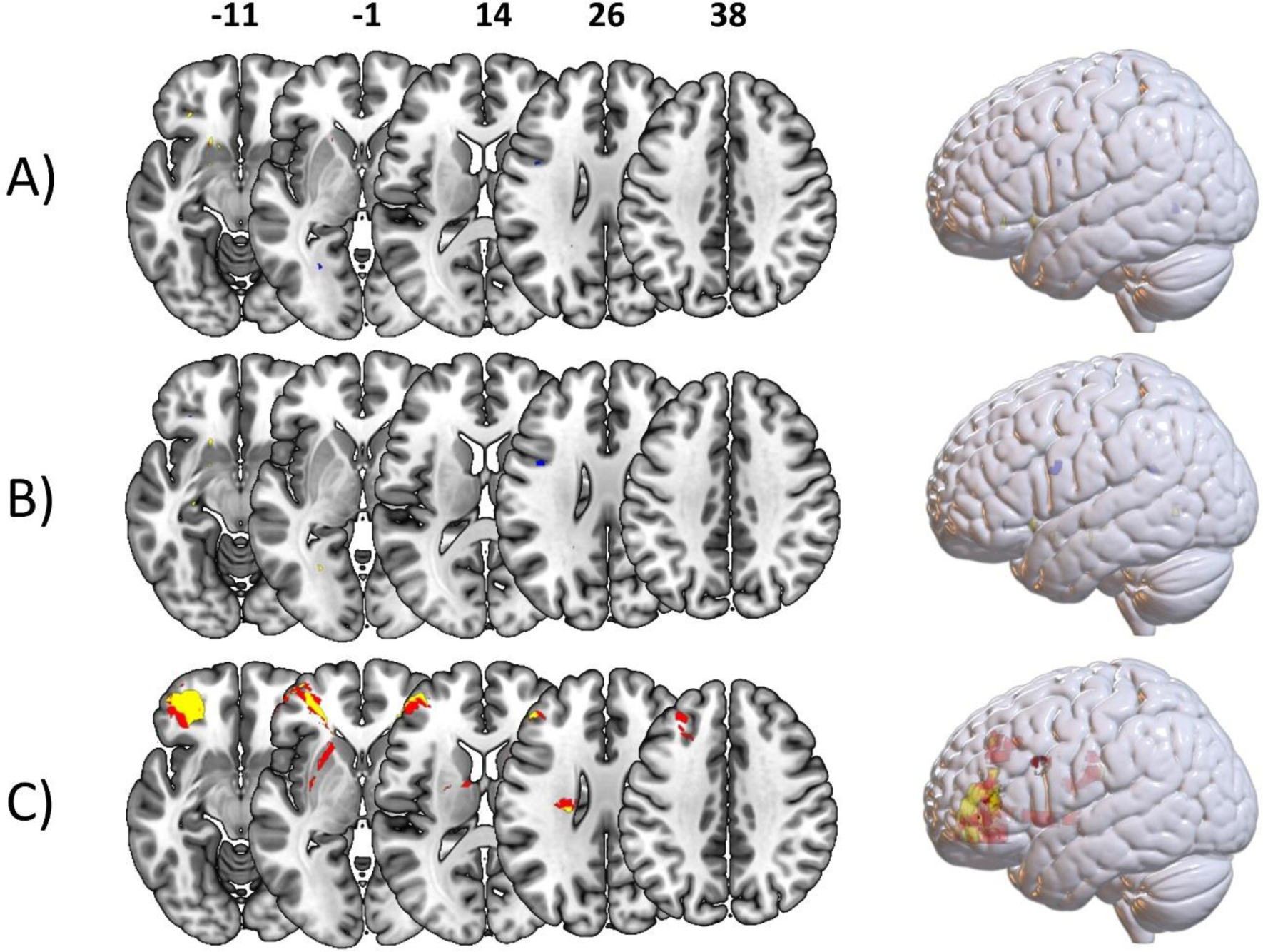
Comparison of results derived from the same analyses with vs. without lesion size control on finger imitation scores. Blue represents voxels with sufficient evidence for the H1 with lesion size control only, red without lesion size control only. Yellow voxels were uncovered by both analyses. Results are displayed from analyses on the A) full sample, B) reduced shared sample, and C) reduced isolated sample.

#### logBFs in relation to z-scores

To have a direct comparison between Bayesian and frequentist statistics, that is logBFs, z-scores and their relation to one another, we created scatterplots for the statistical values resulting from analyses on the the full (see figure 6 A) sample, the reduced shared samples (see figure 6 B) and the reduced isolated sample (see figure 6 C). All in all, the distributions resulting from analyses of both the full and reduced shared sample were skewed towards both lower z-scores and logBFs in reference to the reduced isolated sample. Another noteworthy observation is that a fraction of voxels with sufficient logBFs for an association with hand imitation in the full and shared deficit samples, were found to have positive z-scores. That is, they showed a *reverse association* whereby voxel damage indicated a better performance. When locating these voxels, we found them to form the anterior clusters in the full and reduced shared hand samples (see figure 2 A and B above). In contrast, voxels with strong Bayesian evidence for the H1 in the reduced isolated sample had almost exclusively negative z-scores representing damage leading to worse performance. For a better localization of voxels indicating sufficient evidence for an association, all results were overlapped with anatomical parcellations based on the atlas by Desikan and colleagues (2006). A table containing this information can be found in supplementary table 1.

**Figure 6:**
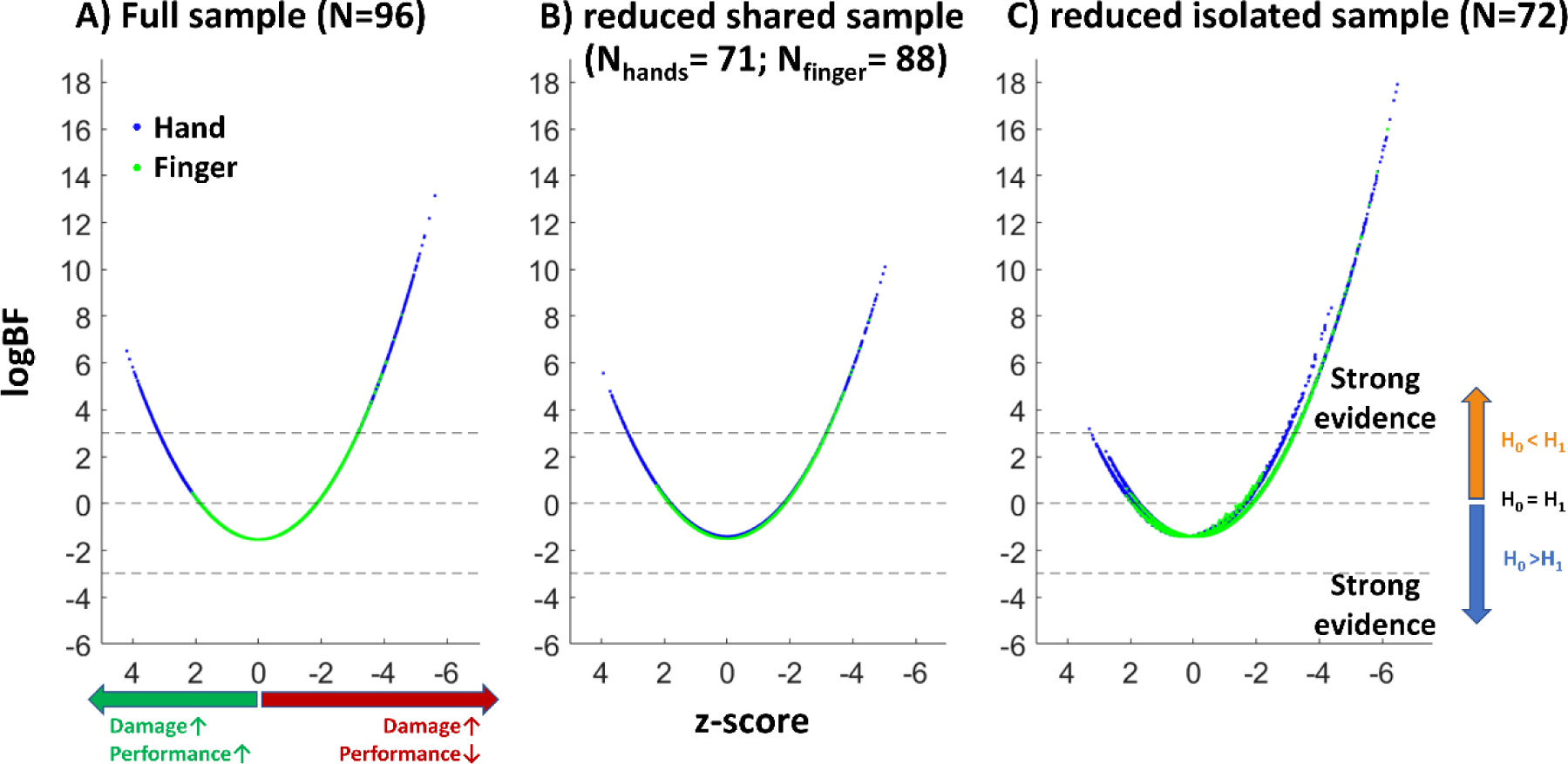
Scatter plots illustrating the relation of logBF and z-score for each analysed voxel. Blue dots represent voxels resulting from analyses concerning hand imitation scores, green dots indicate voxels resulting from analyses concerning finger imitation scores. **A**) contains all voxels from analyses on the full sample. **B**) represents the voxels derived from analyses on the reduced shared samples. **C**) depicts voxels found in analyses in the reduced isolated sample.

### Regionwise lesion-symptom mapping

#### Imitation of hand gestures

As with the VLSM, we report all regions with an evidence BF > 20. In the full sample the regionwise Bayesian lesion-symptom mapping uncovered lesion-symptom associations with the strongest evidence against the null hypothesis between deficits in the imitation of hand gestures and the precuneus (BF=326), followed by the cuneus (BF = 286), superior parietal lobe (BF=143), and the pericalcarine cortex (BF=24). There were no regions surpassing the critical BF in the reduced shared sample. The reduced isolated sample, in contrast, uncovered even more regions and showed overall higher BFs than the full sample. The regions with the strongest evidence against the H0 were the cuneus (BF=5,000,420) and precuneus (BF= 4,822,487), superior parietal cortex (BF=191,644) and the pericalcarine cortex (BF = 2,941), followed by the inferior parietal lobe (BF=54) and paracentral lobule (BF=29). Depictions of the brain regions found in the full and reduced isolated sample can be found in figure 7.

**Figure 7:**
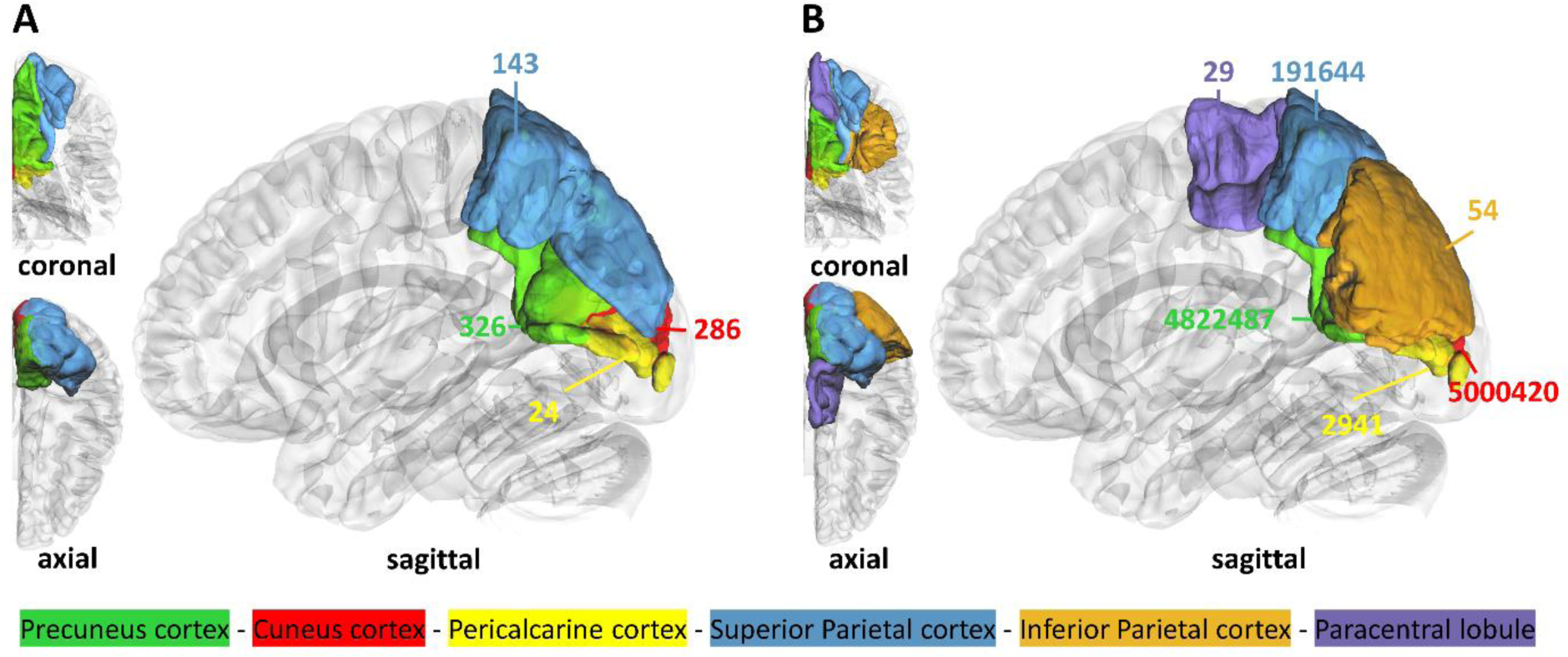
Left hemispheric brain regions with strong evidence against the null hypothesis concerning hand imitation deficits. **A**) Results obtained from the full sample; **B**) regions found in the reduced isolated sample. Numbers next to the coloured regions refer to the corresponding Bayes Factors.

### Imitation of finger gestures

Neither the Bayesian GLMs on the finger imitation scores of the full nor the reduced shared sample uncovered any region with a Bayes factor higher than 20. In the sample where patients with a finger deficit were only included if the deficit was isolated, the rMFG received a BF of 183 (see figure 8).

**Figure 8:**
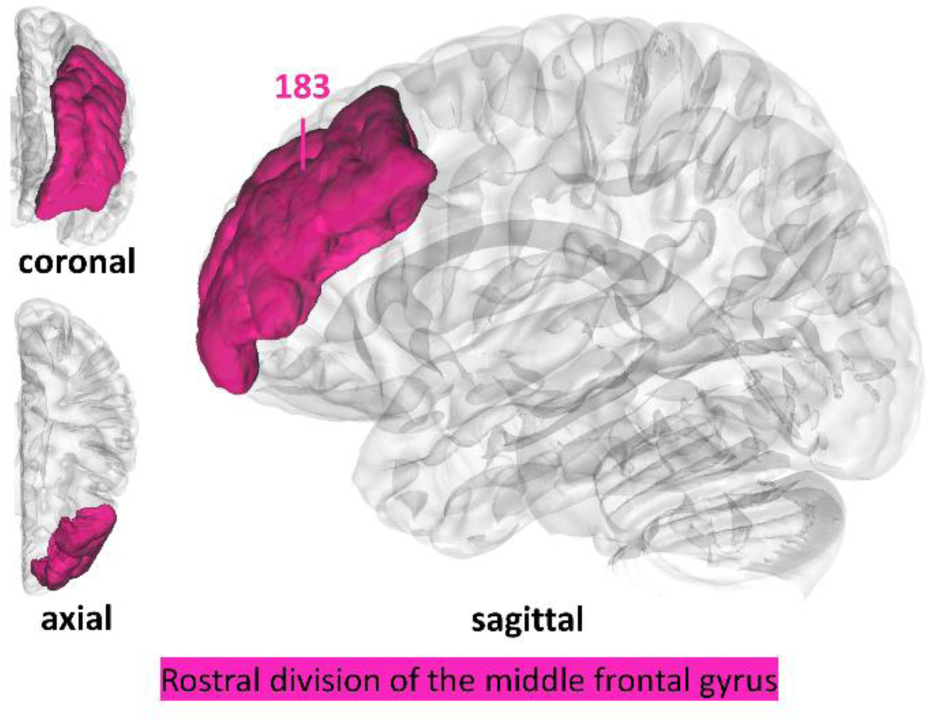
The rMFG in the left hemisphere showed strong evidence for an association with finger imitation deficits in our regionwise analysis testing for Bayesian lesion deficit inference in the reduced isolated sample.

#### logBFs in relation to t-scores

Analyses on patients with exclusively isolated hand deficits boosted both logBFs and t-scores of regions that were found to be associated with a hand deficit in the full sample and uncovered two additional regions with sufficient Bayesian evidence for a region-deficit association. With the exception of the paracentral lobule (Desikan et al., 2007), these regions were also among those with the highest logBFs in the reduced shared sample, albeit not surpassing the critical threshold. rMFG, which had been found in association with isolated finger imitation deficits, persisted around zero regardless of the sample. Interestingly, t-values indicated a trend towards an inverse protective, rather than a damaging effect (for illustrations see figure 9). As can be depicted from the results on the finger imitation scores (figure 10), the rMFG, which demonstrated strong evidence for an association in the reduced isolated sample was assigned a negative logBF in both other samples. On the contrary, regions associated with hand imitation had tendencies towards increased logBFs in the full and reduced shared sample. A summary of all regions, their respective BFs logBFs and t-scores can be found in supplementary table 2 and 3.

**Figure 9:**
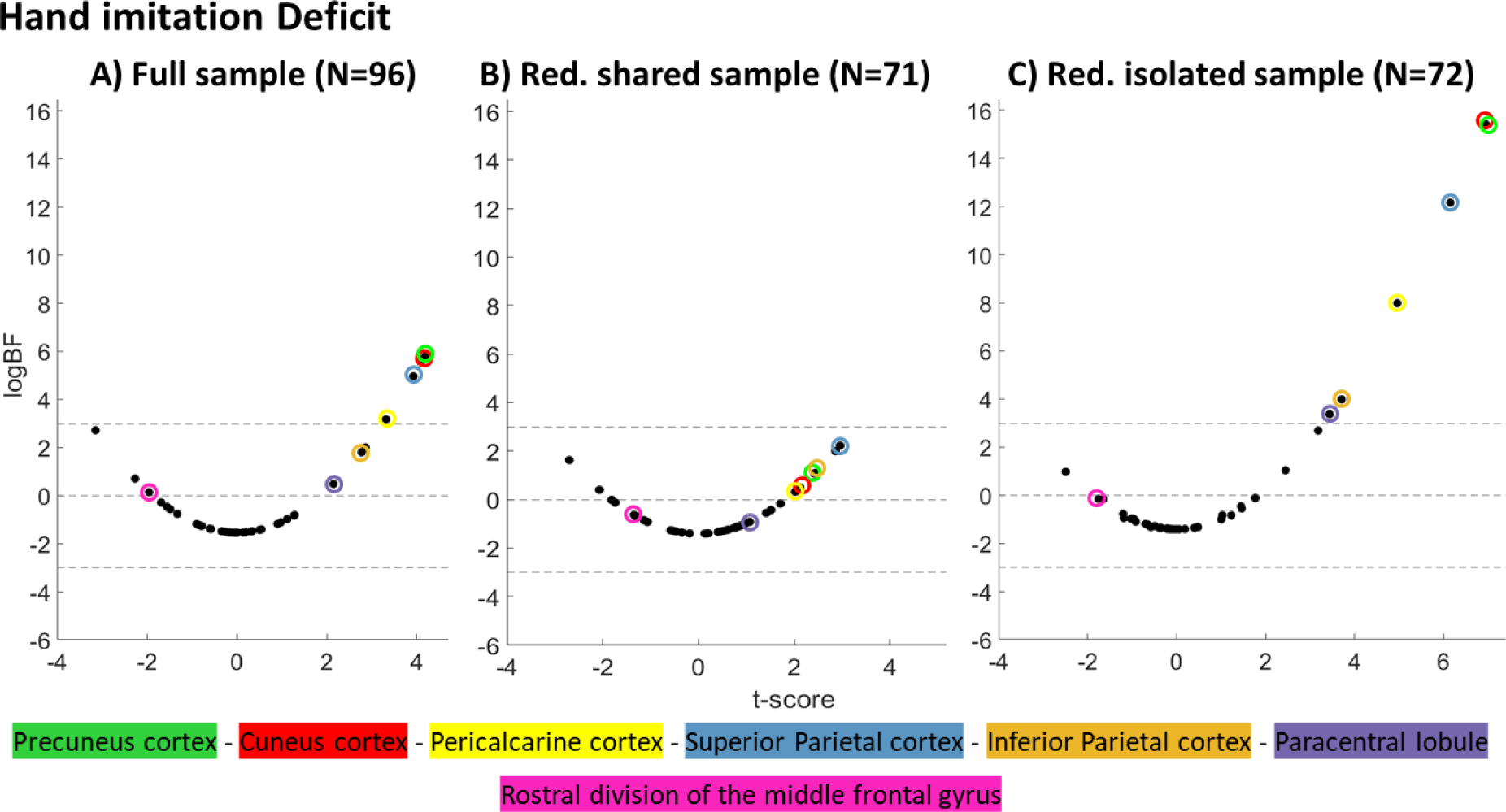
Scatter plots illustrating each region’s logBF relative to its t-score for the three conditions investigating the hand imitation deficit.

**Figure 10:**
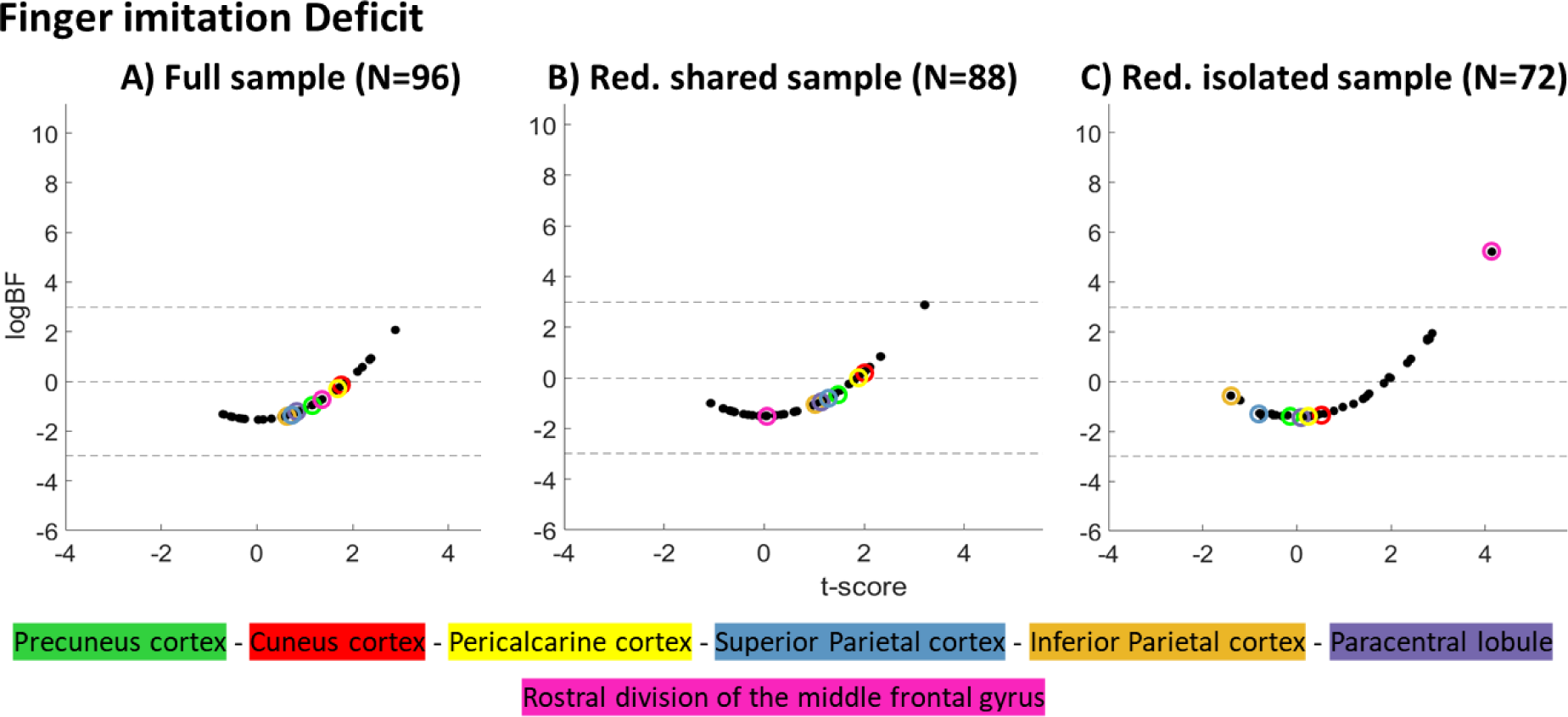
Scatter plots show regionwise logBFs relative to their t-scores for the three samples we investigated in context of deficits in finger imitation.

## Discussion

Our results support the notion of a neural dissociation of finger and hand imitation skills. Hand imitation deficits were found to be associated with damage to more posterior parts of the brain, while isolated finger imitation deficits were located more anteriorly. Isolated hand imitation deficits were associated with damage to mostly parietal and occipital structures. Isolated finger imitation deficits, in contrast, showed a focus in the frontal lobe. While we did uncover one frontal voxel cluster in association with hand imitation deficits, the comparison of frequentist and Bayesian parameters revealed that this cluster had protective rather than damaging properties, as indicated by the positive signs of the frequentist parameters. The dissociation proved stable with and without control for lesion size.

Inferior and superior parietal lobe proved to be the most solid brain-behavior associations concerning hand deficits. They were uncovered by both voxel and regionwise approaches whereby the superior parietal cortex demonstrated both a larger voxel cluster relative to its size (28.34% vs. 22.37%) in the voxelwise lesion-symptom mapping and an overall higher BFs in the regionwise approaches. This is in line with a large proportion of previous research, which reported one or both of these regions as hotspots in context with hand imitation deficits (Achilles et al., 2017; Dovern et al., 2011; Goldenberg & Karnath, 2006; Hoeren et al., 2014). The involvement of precuneus, cuneus, and pericalcarine cortex is, to our knowledge, observed for the first time in the context with hand imitation deficits in lesion symptom mapping. Interestingly, in the regionwise approach these regions showed an even stronger association than the inferior parietal lobe. It might be that cuneus and pericalcarine cortex, in particular, account for proprioceptive aspects of hand imitation deficits. Lesions to these areas have been found in context with deficits in kinesthetic matching, which required the ability to sense the position of once arm in space without visual feedback (Chilvers et al., 2021). It may well be that these regions represent similar proprioceptive deficits in our sample. While our patients were able to see and visually control their finger gestures during the test, hand gestures required the hand to be positioned relative to the face and, therefore, outside the patient’s own visual field, i.e., there was thus no possibility of visual control for hand gestures. Potential proprioceptive deficits could therefore be compensated in the finger but not the hand imitation tasks.

Previous studies stressed, in particular, the inferior frontal gyrus (IFG) and middle frontal gyrus (MFG) in context with finger imitation deficits (Goldenberg & Karnath, 2006; Haaland et al., 2000). Indeed, in our significant voxel cluster in the reduced isolated sample, 18.6 % of the pars orbitalis in the IFG was damaged, which made it the area with the greatest relative damage in this group in the voxelwise approach, followed by rMFG (12.4%). The latter was the only parcel uncovered in the regionwise approach. Thereby, our results located finger imitation even more anteriorly in the IFG than previous studies, which found the center of lesioned mass more in the pars opercularis (Goldenberg & Karnath, 2006). Our results in the middle frontal gyrus closely resemble previous accounts by Haaland and colleagues (2000). They found disproportionally frequent damage to Broadman areas 9 and 46 in patients with impaired finger gestures imitation. Both areas reported by Haaland and colleagues (2000) show considerable overlap with the region defined as rMFG by Desikan and colleagues (2007). Accounts for a frontal lesion focus for finger imitation deficits are not limited to the left, but reported also for the right hemisphere, where the frequency of finger imitation deficits is overall higher (Goldenberg, 1999). To our knowledge, these brain behaviour associations have been less controversial than their counterparts in the left hemisphere. It might be that disconnections to right hemispheric frontal areas via the corpus callosum contribute to isolated finger deficits after left hemispheric frontal damage. When combined with a white matter streamline atlas (Yeh et al., 2018) the rMFG (Desikan et al., 2007) does indeed show a dense structural connection to frontal regions in the opposite hemisphere. This research question requires a more thorough investigation in future works.

Scatterplots illustrating the relation of Bayesian and frequentist lesion mapping results were adopted from the paper by Achilles and colleagues (2017). While the authors limited this illustration to their voxelwise results, we created comparable figures for both voxel- and regionwise brain-behavior associations. They turned out very similar, with the exception that no *reverse associations* surpassed the evidence-threshold in the regionwise Bayesian approaches. The considerably reduced dimensionality of region-wise (as opposed to voxelwise) results allowed the observation of the same regions throughout different analyses. All results support the notion that analyses on the full sample were biased by diluting effects caused by a joint analysis of patients with shared and isolated deficits. The basic principles underlying this mechanism are described by the partial injury problem (cf. Kinkingnéhun et al., 2007; Rorden et al., 2009). In general, regions associated with the relatively larger subgroup (i.e. shared vs. isolated deficit) have a better chance to prevail, while regions from the smaller group will be relatively more watered down, the larger the sample size difference is. Here, the resulting diluted association will be smaller, the smaller the true association of the voxel with the behavior of interest in the sample initially was. Despite a reduced sample size, analyses on our reduced isolated samples consistently uncovered more and overall stronger brain-behaviour associations than those on the full sample containing both shared and isolated imitation deficits.

The inferior parietal cortex and the paracentral lobule were associated with isolated hand deficits but did not persist in our analyses on the full sample. Despite the removal of patients with isolated hand deficits, most isolated hand imitation areas remained among those with the strongest evidence for an association also in the reduced shared sample. Unfortunately, while patients with isolated deficits can be easily distinguished behaviourally, and it seems safe to assume that a patient with an isolated hand deficit will not have suffered damage to a *shared* or *isolated finger* area, the opposite conclusion for a shared deficit cannot be made. There is no easy way to tell if a shared deficit was caused by damage to *shared areas*, damage to both *isolated hand* and *finger areas* or even a mix of *shared* and *isolated areas*. However, high statistical parameters for *isolated hand areas* in the reduced shared sample indicate that lesions leading to a shared deficit often extend to *isolated hand areas*.

Concerning finger imitation deficits, the reduced isolated sample, uncovered associations with the rMFG. In the full sample, consisting of both patients with isolated and shared finger deficits, the rMFG showed a negative logBF, however, not as low as in the reduced shared sample where the rMFG had one of the most negative logBFs. This observation indicates that, unlike *isolated hand areas*, *isolated finger areas* rarely suffer damage in patients with a shared imitation deficit, thereby challenging the notion that simultaneous damage to *isolated hand* and *finger areas* might lead to shared imitation deficits.

In particular patients with a shared imitation deficit should drive the anticipated positive bias for areas truly associated with the respective other imitation deficit, since only their behavioural score can be wrongly associated to areas truly responsible for the respective other deficit. Patients with a shared deficit showed frequent involvement of *isolated hand areas* so that the bias could be nicely demonstrated for *isolated hand areas* in analyses on finger imitation scores. In the reduced isolated sample, patients with a shared deficit were excluded. That is, patients with a finger imitation deficit in this sample did not have accompanying damage to *isolated hand areas*. *Isolated hand areas*, here, were among those with the strongest tendencies against the H1 according to their logBFs or even suggested *reverse associations* according to their t-score. In the full sample, the hypothesized bias was mitigated by the presence of isolated hand deficit patients, who had lesions to the *isolated hand areas* without a finger deficit. In the reduced shared sample, those very patients were excluded, which led the bias to be particularly strong. Cuneus cortex and pericalcarine cortex here even switched sign and became positive (see figure 10 above).

Our results indicate that, in contrast to *isolated hand areas*, *isolated finger areas* were rarely damaged in patients with a shared deficit. For this very reason the rMFG with isolated finger imitation did not bias the analyses on hand imitation skills, as the *isolated hand areas* did with the analyses on finger imitation skills. The bias here would not be expected to begin with, since an area damaged only in the isolated deficit cannot be misattributed to the respective other deficit. Indeed, the logBFs of the rMFG persisted around zero indicating equal evidence for and against the H1 while frequentist parameters indicated a trend towards reverse effects regardless of the sample. This result contests the idea that simultaneous damage to isolated hand and finger areas is responsible for shared imitation deficits.

While we did not uncover any *shared areas,* with sufficient Bayesian evidence, the absence of damage to *isolated finger areas* in patients with a shared deficit indirectly supports their existence. With a BF of 17 also the lingual gyrus was approaching the Bayesian cut-off that we adapted from Achilles and colleagues (2017) in the reduced shared finger sample; it was among the highest ones in the other reduced shared and full samples (ranging from BFs of 7.4 to 7.9), but had considerably lower BFs in the reduced isolated samples (see supplementary table 2). Its posterior location could offer a reasonable explanation why patients with a shared deficit frequently suffered co-occurring damage of *isolated hand areas,* which are in the close proximity to the lingual gyrus, while damage appeared to rarely extend to the more distant *isolated finger areas*. Indeed, the lingual gyrus has previously been found in a functional imaging study, when participants imitated actions from a third-person but not from a first-person perspective (Jackson et al., 2006). The lingual gyrus was therefore suggested to potentially contribute to visuospatial transformation necessary for the former but not for the latter (Jackson et al., 2006). Without the ability to transform observed gestures modelled by someone else onto one’s own body, imitation should fail regardless of the body part to be imitated, which would explain why it affects both hand and finger imitation.

In contrast to Achilles and colleagues (2017), our results did not uncover any voxels surpassing the threshold for evidence for the H0; this might be attributable to our smaller sample size. Bayesian statistics quantify not only evidence for the H1 but also against it. But they do so at different rates, with those in favour of the H1 accumulating considerably faster (Johnson & Rossell, 2010). It has been argued that this asymmetry is unproblematic and simply reflects the fact that support for the absence of something is harder to quantify than support for its presence (van Ravenzwaaij & Wagenmakers, 2022). However, investigating evidence for the H0 will consequently require larger sample sizes than evidence for the H1.

In their regionwise lesion-symptom mapping, Achilles and colleagues (2017) uncovered an almost perfect overlay of regions with sufficient evidence for finger and hand imitation deficits (albeit with variations in the size of the evidence). We did not manage to replicate this result in our full sample, however, based on the effects we uncovered, we are still able to provide two potential explanations how *isolated hand* and *finger areas* might have been covered up in their results. The first one is directly related to our second hypothesis, which assumes a bias introduced by patients with a shared deficit. The second emerged from the dispersions of Bayesian and frequentist statistical parameters in our results. In the following, this is elaborated in more detail.

With respect to the first possible explanation, Bayesian approaches are known to be more liberal than their frequentist counterparts, which is of particular advantage in situations with limited power but, in turn, increases the risk of false positives (Sperber et al., 2023). In a recent simulation study on the voxel level and a sample size closely resembling the one by Achilles and colleagues (2017), the major part of (on average ∼11600) true positives was uncovered, but at the same time one third of all (on average ∼748000) true negatives were attested a wrongful association (Sperber et al., 2023). While these numbers cannot be directly applied on real data, they offer a rough estimate on one of the largest drawbacks of Bayesian approaches. As hypothesized, we could show that an undivided sample with a large number of patients with a shared deficit might produce a systematic bias for regions truly associated with the respective other imitation deficit. Depending on the individual group sizes, *isolated areas* associated with the respective other deficit can even end up with a higher BF than those truly associated to the deficit (cf. figure 10: the cuneus and pericalcarine cortex were assigned a stronger association to finger imitation than the rMFG, which was the only area uncovered in the reduced isolated sample). Apparently overlapping results from an undivided sample, like the one by Achilles and colleagues (2017) might, therefore, represent true associations of the overrepresented imitation type, which also emerged (as false positives) in the other analysis, while the isolated areas of the underrepresented imitation type might be diluted to the point where they are not detected anywhere.

The second possible explanation is that the shared neural correlates for hand and finger imitation in fact represented opposite associations. While Achilles and colleagues (2017) offered insight into the ratios between logBFs and t-scores in their voxelwise approach, regionwise results are reported exclusively for Bayesian analyses. Bayesian statistics offer a quantification of evidence, not only against but also for the H0, however, a clear drawback is that no information can be provided concerning the direction of the uncovered effects. Our VLSMs on hand imitation deficits offer a concrete example of how Bayesian evidence can uncover both the expected negative and reverse positive associations. It is therefore possible that one and the same region is uncovered due to a negative association in one analysis and due to a positive association in the other. Without the combined use of Bayesian and frequentist approaches, there is no way to distinguish them.

In addition to analyses on pure hand and finger imitation scores, Achilles and colleagues (2017) further generated a difference-score for each patient by subtracting the finger imitation score from the hand imitation score. While this measure does indeed estimate the behavioural difference between patients’ hand and finger imitation skills, it is not a sufficient means to provide meaningful brain-behaviour associations which is the reason why we did not (re)analyse this score in the present study. Patients with no deficit at all (e.g. 20-18 points), a deficit in both (e.g. 8-6 points), and an isolated deficit in one imitation type (e.g. 18-16 points), would be assigned an equal difference-score of two, but this similarity lacks any meaningful interpretability on the neural level. Even with a sound sampling approach, results from this analysis cannot be considered as evidence for a task dissociation.

## Conclusion

The present investigations support the notion of a posterior-anterior anatomo-behavioral dissociation for hand and finger imitation skills. It further raises the question whether the lingual gyrus plays a role in (shared) deficits of both hand and finger imitation skills, which should be further investigated in future studies. Beyond, the present analysis demonstrates that wrong sampling can dilute existing associations representing isolated deficits of one imitation skill but not the other and how, in combination with the association problem, it can lead to misleading co-associations between hand and finger imitation deficits. Finally, it shows the exclusive use of Bayesian approaches might uncover opposing associations, which cannot be controlled for. Importantly, both issues can be limited: the former by appropriate sub-sampling, the latter by additional frequentist analyses.

## Acknowledgements

We thank Max Wawrzyniak for providing the MATLAB script for lesion reconstruction.

## Data availability

While public archiving of anonymized patient data is prevented by the ethics approval, access to the data can be provided upon reasonable request by Georg Goldenberg.

## References

Achilles, E. I. S., Weiss, P. H., Fink, G. R., Binder, E., Price, C. J., & Hope, T. M. H. (2017). Using multi-level Bayesian lesion-symptom mapping to probe the body-part-specificity of gesture imitation skills. NeuroImage, 161(January), 94–103. 10.1016/j.neuroimage.2017.08.036

Benjamini, Y., & Hochberg, Y. (1995). Controlling the False Discovery Rate: A Practical and Powerful Approach to Multiple Testing Author (s): Yoav Benjamini and Yosef Hochberg Source: Journal of the Royal Statistical Society. Series B (Methodological), Vol. 57, No. 1 (1995), Publi. Journal of the Royal Statistical Society, 57(1), 289–300.

Bizzozero, I., Costato, D., Della Sala, S., Papagno, C., Spinnler, H., & Venneri, A. (2000). Upper and lower face apraxia: Role of the right hemisphere. Brain, 123(11), 2213–2230. 10.1093/brain/123.11.2213

Chilvers, M. J., Hawe, R. L., Scott, S. H., & Dukelow, S. P. (2021). Investigating the neuroanatomy underlying proprioception using a stroke model. Journal of the Neurological Sciences, 430, 120029. 10.1016/j.jns.2021.120029

De Renzi, E., Motti, F., & Nichelli, P. (1980). Imitating Gestures: A Quantitative Approach to Ideomotor Apraxia. Archives of Neurology, 37(1), 6–10. 10.1001/archneur.1980.00500500036003

Desikan, R. S., Ségonne, F., Fischl, B., Quinn, B. T., Dickerson, B. C., Blacker, D., Buckner, R. L., Dale, A. M., Maguire, R. P., Hyman, B. T., Albert, M. S., & Killiany, R. J. (2006). An automated labeling system for subdividing the human cerebral cortex on MRI scans into gyral based regions of interest. NeuroImage, 31(3), 968–980. 10.1016/j.neuroimage.2006.01.021

Dovern, A., Fink, G. R., Saliger, J., Karbe, H., Koch, I., & Weiss, P. H. (2011). Apraxia impairs intentional retrieval of incidentally acquired motor knowledge. Journal of Neuroscience, 31(22), 8102–8108. 10.1523/JNEUROSCI.6585-10.2011

Goldenberg, G. (1996). Defective imitation of gestures in patients with Damage in the Left or Right Hemispheres. Science, 61(2), 176–180. 10.1136/jnnp.61.2.176

Goldenberg, G. (1999). Matching and imitation of hand and finger postures in patients with damage in the left or right hemispheres. Neuropsychologia, 37(5), 559–566. 10.1016/S0028-3932(98)00111-0

Goldenberg, G. (2003). Pantomime of object use: A challenge to cerebral localization of cognitive function. NeuroImage, 20, 101–106. 10.1016/j.neuroimage.2003.09.006

Goldenberg, G. (2011). Apraxien (A. Thöne-Otto, H. Flor, S. Gauggel, S. Lautenbacher, & H. Niemann (eds.)). Hogrefe.

Goldenberg, G., Hermsdörfer, J., Glindemann, R., Rorden, C., & Karnath, H. O. (2007). Pantomime of tool use depends on integrity of left inferior frontal cortex. Cerebral Cortex, 17(12), 2769–2776. 10.1093/cercor/bhm004

Goldenberg, G., & Karnath, H. O. (2006). The neural basis of imitation is body part specific. The Journal of Neuroscience: The Official Journal of the Society for Neuroscience, 26(23), 6282–6287. 10.1523/JNEUROSCI.0638-06.2006

Goldenberg, G., & Strauss, S. (2002). Hemisphere asymmetries for imitation of novel gestures. Neurology, 59(6), 893–897. 10.1212/WNL.59.6.893

Haaland, K. Y., Harrington, D. L., & Knight, R. T. (2000). Neural representations of skilled movement. Brain, 123(11), 2306–2313. 10.1093/brain/123.11.2306

Hoeren, M., Kümmerer, D., Bormann, T., Beume, L., Ludwig, V. M., Vry, M. S., Mader, I., Rijntjes, M., Kaller, C. P., & Weiller, C. (2014). Neural bases of imitation and pantomime in acute stroke patients: Distinct streams for praxis. Brain, 137(10), 2796– 2810. 10.1093/brain/awu203

Jackson, P. L., Meltzoff, A. N., & Decety, J. (2006). Neural circuits involved in imitation and perspective-taking. NeuroImage, 31(1), 429–439. 10.1016/j.neuroimage.2005.11.026

Johnson, V. E., & Rossell, D. (2010). On the use of non-local prior densities in Bayesian hypothesis tests. Journal of the Royal Statistical Society. Series B: Statistical Methodology, 72(2), 143–170. 10.1111/j.1467-9868.2009.00730.x

Kimura, D., & Archibald, Y. (1974). Motor functions of the left hemisphere. Brain, 97(2), 337–350. 10.1093/brain/97.2.337

Kinkingnéhun, S., Volle, E., Pélégrini-Issac, M., Golmard, J. L., Lehéricy, S., du Boisguéheneuc, F., Zhang-Nunes, S., Sosson, D., Duffau, H., Samson, Y., Levy, R., & Dubois, B. (2007). A novel approach to clinical-radiological correlations: Anatomo-Clinical Overlapping Maps (AnaCOM): Method and validation. NeuroImage, 37(4), 1237–1249. 10.1016/j.neuroimage.2007.06.027

Klingbeil, J., Wawrzyniak, M., Stockert, A., Karnath, H. O., & Saur, D. (2020). Hippocampal diaschisis contributes to anosognosia for hemiplegia: Evidence from lesion network-symptom-mapping. NeuroImage, 208(October 2019), 116485. 10.1016/j.neuroimage.2019.116485

Lausberg, H., & Cruz, R. F. (2004). Hemispheric specialisation for imitation of hand-head positions and finger configurations: A controlled study in patients with complete callosotomy. Neuropsychologia, 42(3), 320–334. 10.1016/j.neuropsychologia.2003.08.003

Lehmkuhl, G., Poeck, K., & Willmes, K. (1982). IDEOMOTOR APRAXIA AND APHASIA: AN EXAMINATION OF Types and Manifestations of apraxic symptoms. Neuropsychologia, 21(3), 199–212. 10.1016/0028-3932(83)90038-6

Liepman, H. (1920). Apraxie. In H. Brugsch (Ed.), Ergebnisse der gesamten Medizin (pp. 516–543). Urban & Schwarzenberg.

Randerath, J. (2023). Syndromes of limb apraxia: Developmental and acquired disorders of skilled movements. In G. G. Brown, T. Z. King, K. Y. Haaland, & B. Crossdon (Eds.), APA handbook of neuropsychology, Vol. 1. Neurobehavioral disorders and conditions: Accepted science and open questions (pp. 159–184). American Psychological Association.

Röhrig, L., Rosenzopf, H., Wöhrstein, S., & Karnath, H. O. (2023). The need for hemispheric separation in pairwise structural disconnection studies. Human Brain Mapping, July, 5212–5220. 10.1002/hbm.26445

Rorden, C., & Brett, M. (2000). Stereotaxic display of brain lesions. Behavioural Neurology, 12(0953–4180), 191–200. www.fil.ion.ucl.ac.uk/spm/

Rorden, C., Fridriksson, J., & Karnath, H. O. (2009). An evaluation of traditional and novel tools for lesion behavior mapping. NeuroImage, 44(4), 1355–1362. 10.1016/j.neuroimage.2008.09.031

Sperber, C., Gallucci, L., Smaczny, S., & Umarova, R. (2023). Bayesian lesion-deficit inference with Bayes factor mapping: key advantages, limitations, and a toolbox. NeuroImage, 271(March), 120008. 10.1016/j.neuroimage.2023.120008

van Ravenzwaaij, D., & Wagenmakers, E. J. (2022). Advantages Masquerading as “Issues” in Bayesian Hypothesis Testing: A Commentary on Tendeiro and Kiers (2019). Psychological Methods, 27(3), 451–465. 10.1037/met0000415

Yeh, F.-C., Panesar, S., Fernandes, D., Meola, A., Masanori;, Y., Fernandez-Miranda, J. C., Vettel, J. M., & Verstynen, T. (2018). Population-Averaged Atlas of the Macroscale Human Structural Connectome and Its Network Topology. NeuroImage, 178, 57–68. doi:10.1016/j.neuroimage.2018.05.027.

