## Supplementary data for "Bayesian evidence for the neural dissociation between finger and hand imitation skills"

### Wrong sampling deludes strong Bayesian evidence for the neural dissociation between finger and hand imitation skills

### Supplementary materials

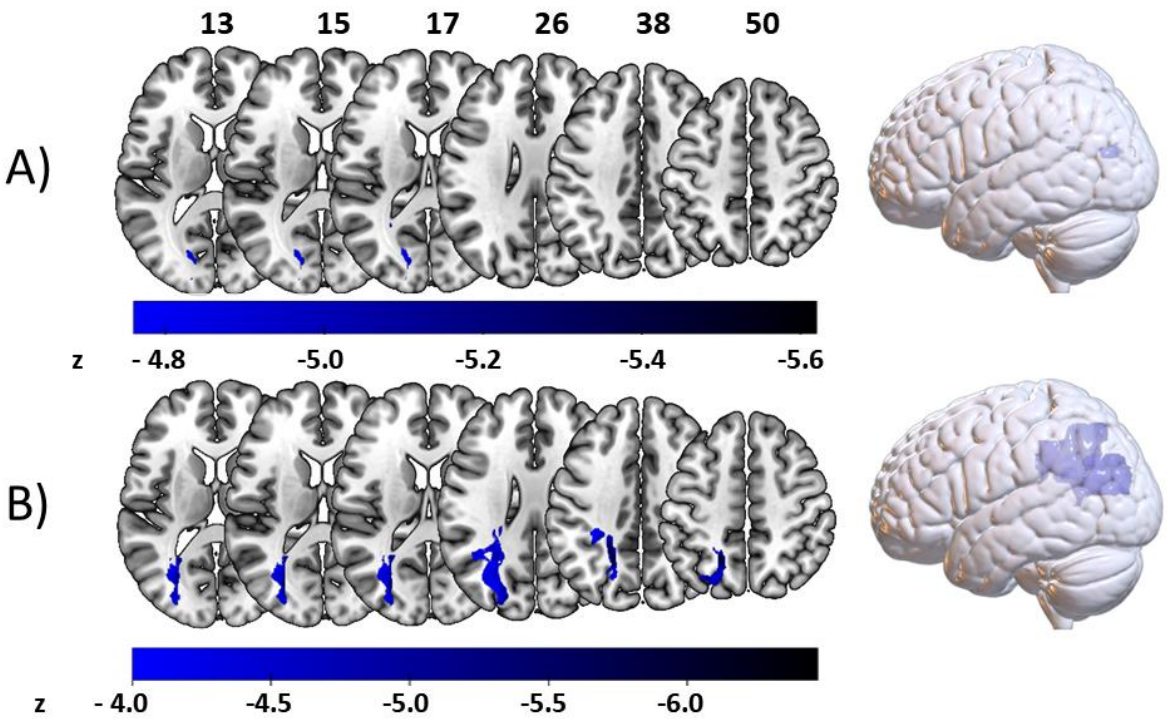


**Supplementary figure 1:** Frequentist associations between voxelwise brain damage and hand imitation deficit. A) Significant voxels resulting from the VLSM on the full sample uncovered a small posterior voxel cluster of 604 voxels ($z_{crit}$=-4.76; $z_{max}$=-5.61). B) The reduced isolated sample uncovered a larger posterior voxel cluster containing 13633 voxels ($z_{crit}$=-4.04; $z_{max}$=-6.47). There were no voxels with sufficient evidence in the reduced shared sample.

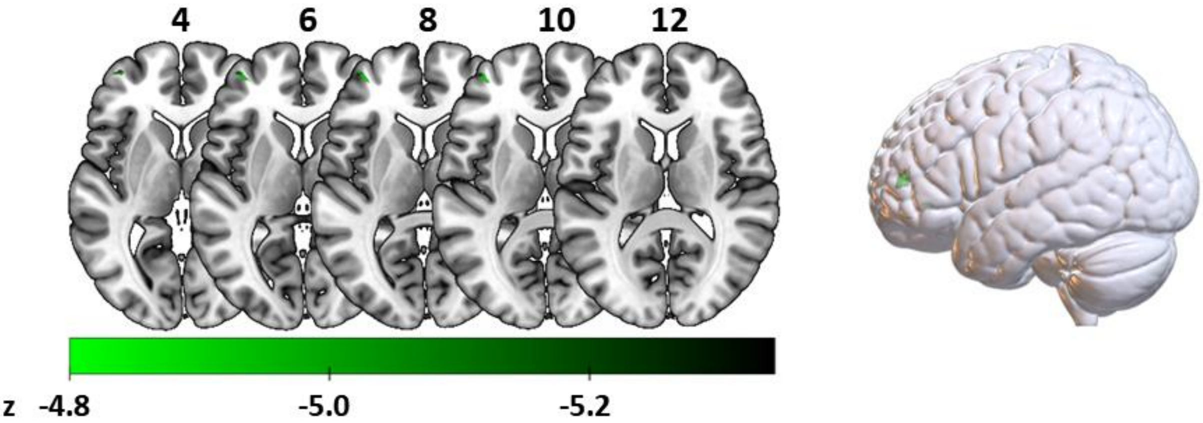


**Supplementary figure 2: The** Frequentist VLSM on finger imitation deficits produced significant results only in the reduced isolated sample, here 353 voxels were significant ($z_{crit}$=-4.83; $z_{max}$=-6.16). Note that for the same VLSM including lesion size control, no significant voxels were found.

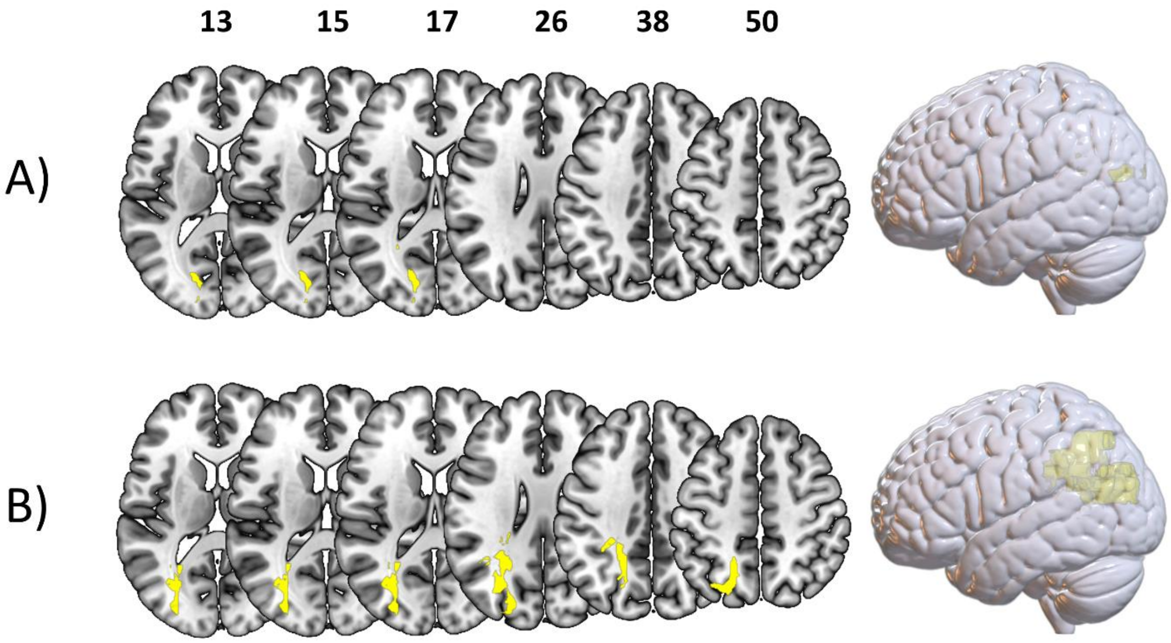


**Supplementary Figure 3:** Comparison of frequentist VLSMs for hand imitation deficits with and without lesion size control. Blue represents voxels with sufficient evidence for the H1 only with lesion size control, red only without lesion size control. Yellow voxels were uncovered by both analyses. A) Shows results of the full sample B) Illustrates voxels found in the reduced isolated sample. Results from both subsamples overlapped almost perfectly.

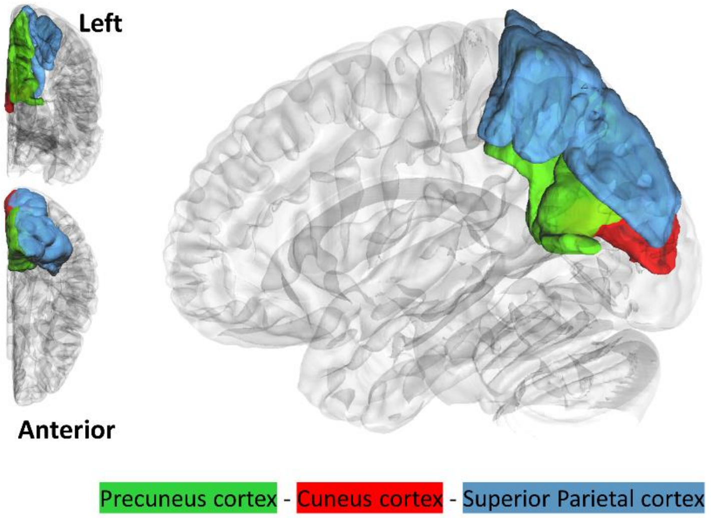


**Supplementary Figure 4**: Frequentist regionwise lesion-symptom mapping, based on the atlas parcellation by Desikan and colleagues (2006) uncovered significant results in only one of the analyses, namely, the analysis of hand imitation skills using the reduced isolated sample. Here, three regions surpassed the critical t-value of 5.64.

**Supplementary Table 1:** Regionwise overlaps of voxels with sufficient Bayesian evidence for results from all sub-samples. abs = absolute numbers of voxels, black cells with white font mark those voxels with sufficient Bayesian evidence, but frequentist parameters indicating a *reverse association*.

| | | Hand all | | Hand shared | | Hand exclusive | | Finger All | | Finger shared | | Finger exclusive | |
|---|---|---|---|---|---|---|---|---|---|---|---|---|---|
| Lobe | Label | abs | % | abs | % | abs | % | abs | % | abs | % | abs | % |
| occipital | Lateraloccipital | 45 | 0.37 | 3 | 0.02 | 19 | 0.16 | 0 | 0.00 | 2 | 0.02 | 0 | 0.00 |
| occipital | Cuneus | 595 | 9.99 | 0 | 0.00 | 0 | 0.00 | 0 | 0.00 | 0 | 0.00 | 0 | 0.00 |
| occipital | Pericalcarine | 336 | 9.29 | 0 | 0.00 | 33 | 0.91 | 0 | 0.00 | 0 | 0.00 | 0 | 0.00 |
| occipital | Lingual | 8 | 0.06 | 0 | 0.00 | 0 | 0.00 | 0 | 0.00 | 19 | 0.15 | 0 | 0.00 |
| parietal | Superiorparietal | 1977 | 10.56 | 267 | 1.43 | 5304 | 28.34 | 0 | 0.00 | 0 | 0.00 | 0 | 0.00 |
| parietal | Inferiorparietal | 1549 | 6.57 | 535 | 2.27 | 5277 | 22.37 | 0 | 0.00 | 0 | 0.00 | 0 | 0.00 |
| parietal | Precuneus | 1831 | 10.31 | 28 | 0.16 | 142 | 0.80 | 0 | 0.00 | 0 | 0.00 | 0 | 0.00 |
| parietal | Supramarginal | 60 | 0.34 | 27 | 0.15 | 2731 | 15.47 | 0 | 0.00 | 0 | 0.00 | 0 | 0.00 |
| parietal | Postcentral | 0 | 0.00 | 0 | 0.00 | 505 | 3.45 | 0 | 0.00 | 0 | 0.00 | 0 | 0.00 |
| temporal | Hippocampus | 0 | 0.00 | 30 | 0.40 | 0 | 0.00 | 0 | 0.00 | 49 | 0.66 | 0 | 0.00 |
| temporal | Banks STS | 8 | 0.15 | 5 | 0.09 | 58 | 1.05 | 0 | 0.00 | 0 | 0.00 | 0 | 0.00 |
| temporal | Superiortemporal | 0 | 0.00 | 0 | 0.00 | 100 | 0.50 | 0 | 0.00 | 0 | 0.00 | 0 | 0.00 |
| insular | Insula | 181 | 1.36 | 21 | 0.16 | 0 | 0.00 | 0 | 0.00 | 6 | 0.05 | 5 | 0.04 |
| subcortical | Amygdala | 0 | 0.00 | 0 | 0.00 | 0 | 0.00 | 1 | 0.03 | 2 | 0.07 | 8 | 0.27 |
| subcortical | Thalamus-Proper | 0 | 0.00 | 0 | 0.00 | 0 | 0.00 | 0 | 0.00 | 0 | 0.00 | 20 | 0.24 |
| subcortical | Accumbens-area | 0 | 0.00 | 0 | 0.00 | 0 | 0.00 | 13 | 1.95 | 0 | 0.00 | 0 | 0.00 |
| subcortical | Caudate' | 0 | 0.00 | 0 | 0.00 | 0 | 0.00 | 33 | 0.57 | 13 | 0.23 | 138 | 2.40 |
| subcortical | Putamen' | 0 | 0.00 | 0 | 0.00 | 0 | 0.00 | 81 | 1.27 | 44 | 0.69 | 512 | 8.00 |
| frontal | Precentral | 798 | 4.14 | 286 | 1.48 | 8 | 0.04 | 0 | 0.00 | 0 | 0.00 | 3 | 0.02 |
| frontal | Caudalmiddlefrontal | 11 | 0.11 | 0 | 0.00 | 0 | 0.00 | 0 | 0.00 | 0 | 0.00 | 6 | 0.06 |
| frontal | Parsopercularis | 787 | 10.30 | 119 | 1.56 | 0 | 0.00 | 0 | 0.00 | 0 | 0.00 | 0 | 0.00 |
| frontal | Parstriangularis | 6 | 0.10 | 0 | 0.00 | 0 | 0.00 | 0 | 0.00 | 0 | 0.00 | 54 | 0.89 |
| frontal | Parsorbitalis | 0 | 0.00 | 0 | 0.00 | 0 | 0.00 | 4 | 0.12 | 0 | 0.00 | 642 | 18.59 |
| frontal | Superiorfrontal | 0 | 0.00 | 0 | 0.00 | 0 | 0.00 | 0 | 0.00 | 0 | 0.00 | 74 | 0.20 |
| frontal | Rostralmiddlefrontal | 14 | 0.06 | 0 | 0.00 | 0 | 0.00 | 8 | 0.03 | 0 | 0.00 | 3053 | 12.42 |
| frontal | Medialorbitofrontal | 0 | 0.00 | 0 | 0.00 | 0 | 0.00 | 5 | 0.06 | 27 | 0.30 | 0 | 0.00 |
| frontal | Lateralorbitofrontal | 0 | 0.00 | 0 | 0.00 | 0 | 0.00 | 51 | 0.39 | 5 | 0.04 | 665 | 5.05 |

**Supplementary table 2:** Brain atlas parcellations as defined by Desikan and colleagues (2007) and their statistical parameters, as uncovered by our GLMs on hand imitation scores on the different subsamples. Bayesian results indicating sufficient evidence/significant frequentist results are presented in bold.

| | **All hand** | | | **shared hand** | | | **exclusive hand** | | |
|---|---|---|---|---|---|---|---|---|---|
| **Label** | **BF** | **logBF** | **t-scores** | **BF** | **logBF** | **t-scores** | **BF** | **logBF** | **t-scores** |
| Thalamus-Proper | 0.28 | -1.27 | -0.78 | 0.28 | -1.29 | -0.53 | 0.25 | -1.40 | -0.14 |
| Caudate | 0.30 | -1.22 | -0.85 | 0.28 | -1.28 | -0.56 | 0.26 | -1.36 | -0.34 |
| Putamen | 0.31 | -1.18 | -0.90 | 0.28 | -1.28 | -0.55 | 0.27 | -1.32 | -0.58 |
| Pallidum | 0.28 | -1.26 | -0.79 | 0.27 | -1.31 | -0.47 | 0.26 | -1.36 | -0.39 |
| Hippocampus | 0.31 | -1.18 | 0.90 | 0.85 | -0.17 | 1.71 | 0.25 | -1.40 | 0.19 |
| Amygdala | 0.22 | -1.53 | -0.10 | 0.26 | -1.34 | 0.41 | 0.24 | -1.41 | -0.15 |
| Accumbens-area | 0.22 | -1.52 | 0.20 | 0.27 | -1.29 | 0.52 | 0.30 | -1.19 | -0.70 |
| Banks STS | 0.37 | -0.99 | 1.12 | 0.57 | -0.55 | 1.42 | 0.43 | -0.83 | 1.23 |
| Caudalanteriorcingulate | 0.22 | -1.53 | -0.16 | 0.25 | -1.40 | 0.13 | 0.46 | -0.77 | -1.20 |
| Caudalmiddlefrontal | 0.75 | -0.29 | -1.69 | 0.99 | -0.01 | -1.81 | 0.28 | -1.28 | -0.49 |
| Cuneus | **286.12** | **5.66** | 4.14 | 1.63 | 0.49 | 2.12 | **5000420.36** | **15.43** | **6.95** |
| Entorhinal | 0.21 | -1.54 | -0.02 | 0.33 | -1.10 | 0.85 | 0.24 | -1.41 | -0.01 |
| Fusiform | 0.22 | -1.53 | 0.13 | 0.32 | -1.14 | 0.78 | 0.26 | -1.35 | 0.40 |
| Inferiorparietal | 6.04 | 1.80 | 2.78 | 3.01 | 1.10 | 2.45 | **53.65** | **3.98** | 3.71 |
| Inferiortemporal | 0.21 | -1.54 | 0.01 | 0.31 | -1.17 | 0.74 | 0.24 | -1.41 | 0.05 |
| Isthmuscingulate | 0.44 | -0.81 | 1.28 | 0.00 | - | 0.00 | 0.89 | -0.12 | 1.77 |
| Lateraloccipital | 0.25 | -1.41 | 0.55 | 0.28 | -1.26 | 0.59 | 0.58 | -0.55 | 1.46 |
| Lateralorbitofrontal | 0.25 | -1.39 | -0.58 | 0.25 | -1.39 | -0.18 | 0.37 | -0.98 | -1.02 |
| Lingual | 7.39 | 2.00 | 2.86 | 7.37 | 2.00 | 2.86 | 2.81 | 1.03 | 2.45 |
| Medialorbitofrontal | 0.24 | -1.43 | 0.50 | 0.32 | -1.14 | 0.79 | 0.33 | -1.10 | -0.92 |
| Middletemporal | 0.22 | -1.52 | -0.20 | 0.27 | -1.32 | 0.45 | 0.25 | -1.38 | -0.21 |
| Parahippocampal | 0.22 | -1.49 | 0.32 | 0.65 | -0.43 | 1.52 | 0.27 | -1.32 | 0.48 |
| Paracentral | 1.62 | 0.48 | 2.15 | 0.40 | -0.92 | 1.07 | **29.04** | **3.37** | 3.44 |
| Parsopercularis | 15.15 | 2.72 | -3.15 | 5.10 | 1.63 | -2.70 | 2.65 | 0.98 | -2.49 |
| Parsorbitalis | 0.47 | -0.76 | -1.33 | 0.50 | -0.68 | -1.30 | 0.30 | -1.21 | -0.66 |
| Parstriangularis | 2.02 | 0.70 | -2.27 | 0.88 | -0.12 | -1.74 | 0.85 | -0.16 | -1.75 |
| Pericalcarine | **23.90** | **3.17** | 3.32 | 1.38 | 0.32 | 2.02 | **2941.26** | **7.99** | 4.96 |
| Postcentral | 0.23 | -1.49 | -0.34 | 0.40 | -0.92 | -1.07 | 0.63 | -0.46 | 1.45 |
| Posteriorcingulate | 0.33 | -1.12 | 0.98 | 0.36 | -1.01 | 0.97 | 0.36 | -1.01 | 1.00 |
| Precentral | 0.62 | -0.47 | -1.56 | 1.49 | 0.40 | -2.07 | 0.25 | -1.40 | -0.21 |
| Precuneus | **326.45** | **5.79** | 4.18 | 2.90 | 1.06 | 2.43 | **4822486.78** | **15.39** | **6.96** |
| Rostralanteriorcingulate | 0.21 | -1.54 | -0.06 | 0.25 | -1.39 | 0.20 | 0.38 | -0.97 | -0.99 |
| Rostralmiddlefrontal | 1.15 | 0.14 | -1.96 | 0.53 | -0.64 | -1.35 | 0.85 | -0.16 | -1.65 |
| Superiorfrontal | 0.25 | -1.38 | -0.60 | 0.26 | -1.36 | -0.35 | 0.37 | -1.00 | -0.95 |
| Superiorparietal | **143.13** | **4.96** | 3.93 | 9.21 | 2.22 | 2.96 | **191643.60** | **12.16** | **6.15** |
| Superiortemporal | 0.22 | -1.51 | -0.27 | 0.25 | -1.40 | 0.17 | 0.24 | -1.41 | -0.08 |
| Supramarginal | 0.44 | -0.82 | 1.28 | 0.29 | -1.24 | 0.62 | 14.71 | 2.69 | 3.18 |
| Temporalpole | 0.23 | -1.49 | 0.34 | 0.34 | -1.06 | 0.90 | 0.25 | -1.40 | -0.17 |
| Transversetemporal | 0.22 | -1.52 | -0.18 | 0.28 | -1.26 | -0.58 | 0.44 | -0.83 | 1.03 |
| Insula | 0.57 | -0.57 | -1.48 | 0.43 | -0.85 | -1.14 | 0.39 | -0.95 | -1.19 |

**Supplementary table 3:** Brain atlas parcellations as defined by Desikan and colleagues (2007) and their statistical parameters, as uncovered by our GLMs on finger imitation scores on the different subsamples. Bayesian results indicating sufficient evidence/significant frequentist results are presented in bold.

| | All finger | | | shared finger | | | exclusive finger | | |
|---|---|---|---|---|---|---|---|---|---|
| **Label** | **BF** | **logBF** | **t-scores** | **BF** | **logBF** | **t-scores** | **BF** | **logBF** | **t-scores** |
| Thalamus-Proper | 0.22 | -1.51 | -0.26 | 0.23 | -1.48 | -0.23 | 0.25 | -1.39 | 0.27 |
| Caudate | 0.30 | -1.19 | 0.88 | 0.24 | -1.43 | 0.40 | 1.16 | 0.15 | 1.99 |
| Putamen | 2.41 | 0.88 | 2.35 | 0.91 | -0.10 | 1.80 | 5.53 | 1.71 | 2.82 |
| Pallidum | 0.39 | -0.94 | 1.16 | 0.33 | -1.10 | 0.95 | 0.61 | -0.49 | 1.53 |
| Hippocampus | 1.49 | 0.40 | 2.10 | 2.33 | 0.84 | 2.33 | 0.26 | -1.35 | 0.37 |
| Amygdala | 0.94 | -0.07 | 1.83 | 1.06 | 0.05 | 1.89 | 0.31 | -1.18 | 0.79 |
| Accumbens-area | 0.74 | -0.31 | 1.67 | 0.61 | -0.49 | 1.52 | 0.36 | -1.02 | 0.98 |
| Banks STS | 0.22 | -1.49 | -0.32 | 0.22 | -1.50 | -0.10 | 0.28 | -1.29 | -0.71 |
| Caudalanteriorcingulate | 1.49 | 0.40 | 2.10 | 0.51 | -0.67 | 1.38 | 5.59 | 1.72 | 2.77 |
| Caudalmiddlefrontal | 0.21 | -1.54 | 0.03 | 0.27 | -1.31 | -0.66 | 0.94 | -0.07 | 1.85 |
| Cuneus | 0.96 | -0.04 | 1.85 | 1.45 | 0.37 | 2.08 | 0.27 | -1.31 | 0.47 |
| Entorhinal | 0.30 | -1.21 | 0.86 | 0.33 | -1.10 | 0.96 | 0.25 | -1.37 | 0.35 |
| Fusiform | 0.33 | -1.11 | 0.98 | 0.42 | -0.87 | 1.20 | 0.24 | -1.41 | 0.13 |
| Inferiorparietal | 0.25 | -1.37 | 0.62 | 0.39 | -0.94 | 1.14 | 0.57 | -0.57 | -1.40 |
| Inferiortemporal | 0.27 | -1.30 | 0.73 | 0.35 | -1.04 | 1.02 | 0.25 | -1.38 | -0.32 |
| Isthmuscingulate | 0.25 | -1.41 | -0.55 | 0.24 | -1.43 | -0.41 | 0.25 | -1.40 | -0.72 |
| Lateraloccipital | 0.46 | -0.78 | 1.31 | 0.78 | -0.24 | 1.70 | 0.47 | -0.75 | -1.20 |
| Lateralorbitofrontal | 2.54 | 0.93 | 2.38 | 1.25 | 0.22 | 1.99 | 2.48 | 0.91 | 2.42 |
| Lingual | 7.95 | 2.07 | 2.89 | 17.77 | 2.88 | 3.21 | 0.26 | -1.35 | -0.45 |
| Medialorbitofrontal | 1.79 | 0.58 | 2.20 | 1.53 | 0.43 | 2.11 | 0.25 | -1.37 | 0.36 |
| Middletemporal | 0.22 | -1.50 | 0.30 | 0.26 | -1.34 | 0.60 | 0.28 | -1.29 | -0.54 |
| Parahippocampal | 0.22 | -1.53 | 0.13 | 0.23 | -1.48 | 0.23 | 0.25 | -1.39 | 0.26 |
| Paracentral | 0.29 | -1.24 | 0.82 | 0.34 | -1.08 | 0.98 | 0.24 | -1.41 | 0.10 |
| Parsopercularis | 0.24 | -1.42 | -0.51 | 0.37 | -1.00 | -1.07 | 0.41 | -0.90 | 1.20 |
| Parsorbitalis | 0.75 | -0.29 | 1.69 | 0.35 | -1.05 | 1.01 | 5.24 | 1.66 | 2.77 |
| Parstriangularis | 0.28 | -1.29 | 0.75 | 0.22 | -1.50 | 0.05 | 2.11 | 0.75 | 2.35 |
| Pericalcarine | 0.80 | -0.22 | 1.73 | 1.26 | 0.23 | 2.00 | 0.25 | -1.40 | 0.24 |
| Postcentral | 0.26 | -1.33 | -0.69 | 0.27 | -1.29 | -0.68 | 0.28 | -1.28 | 0.59 |
| Posteriorcingulate | 0.68 | -0.39 | 1.62 | 0.91 | -0.09 | 1.80 | 0.25 | -1.38 | 0.31 |
| Precentral | 0.23 | -1.48 | -0.37 | 0.30 | -1.21 | -0.82 | 0.54 | -0.61 | 1.47 |
| Precuneus | 0.39 | -0.95 | 1.15 | 0.54 | -0.62 | 1.42 | 0.26 | -1.35 | -0.20 |
| Rostralanteriorcingulate | 0.98 | -0.02 | 1.86 | 0.39 | -0.94 | 1.13 | 1.19 | 0.17 | 1.96 |
| Rostralmiddlefrontal | 0.49 | -0.72 | 1.36 | 0.22 | -1.50 | 0.03 | **183.65** | **5.21** | 4.14 |
| Superiorfrontal | 0.74 | -0.30 | 1.68 | 0.27 | -1.32 | 0.63 | 6.94 | 1.94 | 2.87 |
| Superiorparietal | 0.27 | -1.31 | 0.72 | 0.41 | -0.89 | 1.18 | 0.28 | -1.27 | -0.79 |
| Superiortemporal | 0.24 | -1.42 | 0.52 | 0.27 | -1.31 | 0.65 | 0.24 | -1.41 | 0.13 |
| Supramarginal | 0.27 | -1.32 | -0.71 | 0.26 | -1.35 | -0.59 | 0.26 | -1.35 | -0.50 |
| Temporalpole | 0.90 | -0.11 | 1.80 | 1.30 | 0.26 | 2.02 | 0.25 | -1.40 | 0.22 |
| Transversetemporal | 0.22 | -1.50 | -0.29 | 0.23 | -1.46 | -0.32 | 0.26 | -1.36 | 0.42 |
| Insula | 0.25 | -1.37 | 0.62 | 0.23 | -1.45 | 0.33 | 0.50 | -0.70 | 1.41 |
